# TeTIR domains define a new class of animal TIR-like NADase effectors in STAND proteins

**DOI:** 10.64898/2026.09.21.753143

**Authors:** Joelle Hornebeck, Ruslan Ibragimov, Nicole Oelerich, Paolo Pfad, Jan Jirschitzka, Jan Gebauer, Ulrich Baumann, Thomas Hermanns, Kay Hofmann

**Affiliations:** Institute for Genetics, University of Cologne, Zülpicher Straße 47a, D-50674 Cologne, Germany; Institute of Biochemistry, University of Cologne, Zülpicher Straße 47, D-50674 Cologne, Germany

## Abstract

TIR domains are found in diverse immune signaling proteins and, in several systems, function as NAD+-cleaving enzymes causing or signaling cell death. In plants, TIR domains occur as effector domains in TNL immune receptors, where they are coupled to a central STAND ATPase and a C-terminal repeat domain. Here, we identify TeTIR (TEP1 extended TIR), a previously unrecognized family of TIR-like domains associated with NACHT-type ATPases and C-terminal WD40 or TPR repeat domains. The TeTIR domain of human TEP1 exhibits robust NADase activity comparable to SARM1-TIR, making TEP1 the only other NADase-active TIR-like protein identified in mammals. NADase activity is conserved in TEP1 homologs from diverse eukaryotes. The crystal structure of TEP1 TeTIR reveals an extended TIR-like fold with additional secondary-structure elements and a distinctive RWG-containing loop. Molecular dynamics simulations indicate that the catalytic glutamate contributes to catalysis and the conformational equilibrium of the active-site region, promoting formation of a substrate-accessible state. Structure-guided mutagenesis identifies conserved residues contributing to NAD+ recognition and catalysis, including features distinct from canonical TIR NADases. Expression of isolated TEP1 TeTIR causes NAD+ depletion and cell death in a catalytic-glutamate-dependent manner. TeTIRs thus define a distinct family of animal TIR-like NADases embedded in NACHT-type STAND architectures.

## Introduction

TIR (Toll/interleukin-1 receptor) domains were first identified in the intracellular regions of Toll and interleukin-1 receptors and were initially considered to mediate protein–protein interactions in innate immune signaling at the membrane ^1^. TIR domains can form homo- and heterotypic assemblies, ranging from dimers and trimers to higher-order filament-like structures, thereby providing a signaling platform through regulated oligomerization. In this respect, TIR domains functionally resemble the death-fold domains (DD, death domain; DED, death effector domain; CARD, caspase recruitment domain; and PYD, pyrin domain), which likewise mediate signaling through homotypic interactions and higher-order assembly, although the two domain families are structurally unrelated ^2^. In animals, TIR domains are predominantly found in the cytoplasmic regions of Toll-like and interleukin-1 receptor family proteins and their adaptor proteins. In plants, by contrast, TIR domains are predominantly found at the N-termini of intracellular immune receptors of the TNL class. These proteins combine an N-terminal TIR effector domain with a central NB-ARC-type ATPase and a C-terminal leucine-rich repeat (LRR) region ^3, 4^. Recognition of pathogen-derived effectors promotes oligomerization of the NB-ARC domain and assembly of receptor complexes termed resistosomes, activating the TIR domains through induced proximity. This architecture has striking parallels to the mammalian inflammasome and apoptosome, in which ligand-induced oligomerization of STAND ATPase domains also generates higher-order signaling platforms bearing N-terminal effector domains such as CARD or PYD domains.

The underlying architectures nevertheless differ: plant TNLs employ an NB-ARC ATPase and LRR sensor domain, whereas canonical mammalian inflammasome NLRs contain NACHT ATPases, and the apoptosome-forming protein APAF1 combines an NB-ARC domain with C-terminal WD40 repeats. Both NB-ARC and NACHT ATPases belong to the STAND (signal transduction ATPases with numerous domains) superfamily. Thus, plant resistosomes, inflammasomes and apoptosomes represent evolutionary distinct versions of a common organizational principle in which a sensory repeat domain controls oligomerization of a central STAND ATPase, thereby activating an N-terminal effector domain ^5^.

Unlike their mammalian counterparts, plant TIR domains within immune receptors do not signal through recruitment of TIR-containing adaptor proteins, but rather through the enzymatic activity of the TIR domains themselves. Plant TIRs hydrolyze NAD^+^ to generate nicotinamide (NAM) and ADP-ribose (ADPR), and can further convert these products into a diverse set of cyclic and ADP-ribosylated nucleotides ^6, 7^. These products include ADP-ribosylated ATP (ADPr-ATP) and di-ADPR, which promote cell death via the EDS1/SAG101–NRG1 axis ^8^. Bacteria also have TIR domains with NADase activity involved in cell death, typically as part of the antiphage defense. In the Pycsar system, the NADase activity of the PycTIR protein is activated by cUMP or cCMP, leading to the cleavage of NAD^+^ to NAM and ADPR and, ultimately, cell death through NAD^+^ depletion ^9^. Similarly, bacterial CBASS-associated TIR-SAVED systems are activated by cyclic oligonucleotides, triggering higher-order TIR filament formation and NAD^+^ depletion, resulting in abortive infection and cell death ^10^. In contrast to plant TIRs, these bacterial cell-death mechanisms do not appear to depend on NAD-derived metabolites.

In mammals, most TIR domains are considered enzymatically inactive, either through loss of active site residues or due to structural features incompatible with catalysis. The only established exception is SARM1, a TIR-domain protein that is neither a receptor nor an adaptor ^11^. SARM1 plays a central role in Wallerian degeneration following axotomy, in which disruption of axonal transport leads to the loss of the short-lived NAD^+^-biosynthetic enzyme NMNAT2, resulting in decreased NAD^+^ synthesis and accumulation of its precursor NMN ^12^. SARM1 is normally kept inactive by NAD^+^ binding to its autoinhibitory ARM domain ^13^. An increase in the NMN/NAD^+^ ratio relieves this inhibition, allowing the TIR domains to oligomerize and activate their NADase activity ^14^, thereby driving a rapid depletion of NAD^+^ in the severed axon to levels insufficient for axonal survival ^15^.

Here, we identify a previously unrecognized family of TIR-like domains in proteins with a TNL-like architecture, combining an N-terminal TIR-like effector domain with a central NACHT-type STAND ATPase and C-terminal WD40 repeats. By systematic bioinformatic analysis of N-terminal regions associated with STAND ATPases from diverse organisms, we identified a conserved family of TIR-like domains, including one copy in human TEP1. We termed these domains TeTIR (TEP1 extended TIR) based on their remote sequence similarity to TIR domains and their substantially larger size resulting from several insertions. Sequence and structural analyses revealed conserved features characteristic of an active TIR-like NADase, including a conserved active site glutamate residue. We characterized the TeTIR domain of human TEP1 and found that it possesses robust NADase activity comparable to that of SARM1. Expression of the isolated TeTIR domain in cells resulted in pronounced NAD^+^ depletion and cell death, suggesting that TeTIR domains represent a previously unrecognized class of animal TIR-like NADases capable of coupling NAD^+^ depletion to cell death.

## Results

### Identification of the TeTIR domain family

STAND-type ATPases from animals, plants, fungi, and bacteria frequently adopt a tripartite architecture, in which an N-terminal effector domain precedes the NACHT- or NB-ARC-type ATPase domain. To explore the repertoire of effector domains associated with STAND ATPases across different taxa, we performed a bioinformatic analysis of their N-terminal regions. While many mammalian STAND ATPases harbor N-terminal CARD or PYD domains, we identified a previously uncharacterized family of NACHT ATPases carrying a different N-terminal domain. The family comprises five proteins encoded in the human genome, including TEP1, NWD1, and NWD2, as well as related proteins found in other animals, a small number of fungi and protists, and several bacteria and archaea. No representatives were identified in plants. The domain comprises 200–350 residues and is invariably found directly upstream of the NACHT domain. In most proteins, it forms the N-terminal region, while TEP1 and its orthologs are N-terminally extended by a TROVE (Telomerase, Ro, Vault) and VWA (von Willebrand factor type A) domain pair. The C-terminal part of the proteins contains either WD40 or TPR-type repeat regions (Figure 1A, Supp. Fig 1A).

**Figure 1:**
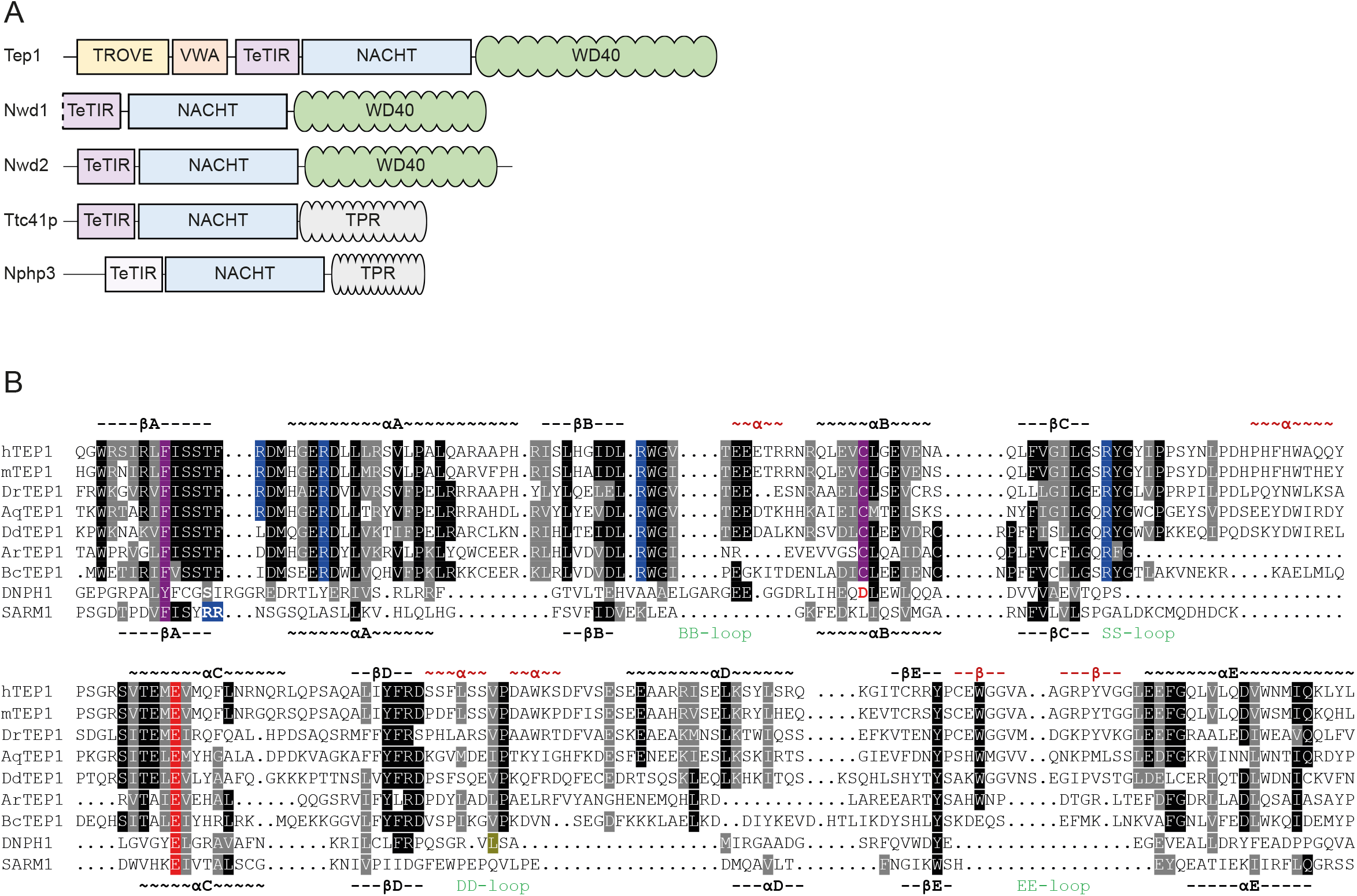
The TeTIR family. (**A**) Domain architecture of human TeTIR family members, shown approximately to scale. TeTIR domains (purple) and NACHT-type ATPase domains (blue) are present in all proteins; the catalytically inactive TeTIR domain of NPHP3 is shown in light purple. Other domains are indicated by colored boxes. (**B**) Multiple sequence alignment of representative TeTIR domains, alongside the TIR domain of SARM1 and the catalytic domain of DNPH1. Residues invariant or conservatively substituted in ≥50% of the sequences are shown on black or grey backgrounds, respectively. Functional residues analyzed in this study are highlighted. Secondary structure elements of TEP1-TeTIR and SARM1-TIR are shown above and below the alignment, respectively. Red elements indicate TeTIR-specific insertions relative to the core TIR fold.

Most non-mammalian members of the family have a TEP1-like architecture, although the prokaryotic proteins lack the N-terminal extension, and their relationship to human TEP1 and to each other could be established by sequence-to-sequence comparison. The other mammalian proteins (NWD1, NWD2, TTC41, and NPHP3) form more divergent subfamilies, whose relationship to TEP1 was established using the generalized profile method ^16^. Pairwise comparisons between the different subfamilies yielded significant p-values in all cases. We therefore combined all subfamilies into a common multiple alignment, which was subsequently compared with databases of known protein and domain alignments using hidden Markov model (HMM)–to–HMM comparison ^17^.

In these searches, we detected a distant but statistically significant relationship to HMMs derived from TIR-domain alignments, as well as to those derived from MilB/DNPH family alignments. TIR-type NADases and MilB/DNPH enzymes share a common catalytic mechanism for hydrolytically cleaving N-glycosidic bonds between nitrogenous bases and phosphoribose ^18, 19^. Moreover, these enzyme families are structurally related and share a conserved catalytic glutamate residue ^14^. Strikingly, a glutamate corresponding to the conserved catalytic residue is present in TEP1, NWD1, NWD2, and TTC41, but is absent from NPHP3, suggesting that members of the domain family may possess NADase activity. Structure prediction using AlphaFold ^20^ further supported a TIR-like fold, with several additional loops extending from the core fold. We therefore refer to this previously unrecognized domain family as TeTIR (<u>T</u>EP1 <u>e</u>xtended <u>TIR</u>), an alignment of representative members is shown in Figure 1B. The AlphaFold model of NWD1 suggested that its TeTIR domain is N-terminally truncated, resulting in the loss of an internal β-strand present in the closely related NWD2 and all other family members. Human TTC41 is a pseudogene (TTC41P) and is therefore unlikely to encode a functional protein. Since NPHP3 lacks the putative catalytic glutamate, TEP1 and NWD2 were the only plausible candidates among the human proteins for encoding active enzymes.

### TEP1 TeTIR is an NADase

To investigate the predicted NADase activity, we expressed and purified recombinant human TEP1 TeTIR domain (residues 895–1117) from E. coli. Following affinity purification and removal of the purification tag, the TEP1 TeTIR domain eluted as a single, well-defined species by size-exclusion chromatography, consistent with a monomeric state (Suppl. Fig. S2A). Purified human TEP1 TeTIR was then incubated with NAD^+^ and the reaction was monitored by targeted LC–MS/MS. Canonical TIR domains have been shown to produce not only nicotinamide (NAM) and ADPR, but also several ADPR derivatives, including cADPR, 2⍰ADPR, and 3⍰ADPR. These products were therefore monitored alongside NAM and ADPR, with product identities assigned based on their mass-to-charge ratios and chromatographic retention times relative to authentic standards.

Incubation of TEP1 TeTIR with NAD^+^ resulted in a time-dependent decrease in NAD^+^ over 6 h, accompanied by the accumulation of NAM (Fig. 2A), thus demonstrating NADase activity of the isolated domain. To assess the dependence of this activity on the predicted catalytic glutamate, we next analyzed NAM and ADPR generation by wild-type and glutamate-mutant TeTIR domains from human and mouse TEP1. Both hTEP1 and mTEP1 generated NAM and ADPR efficiently, whereas substitution of the predicted catalytic glutamate (E1008A in hTEP1 and E1017A in mTEP1) abolished product formation, with levels comparable to those in enzyme-free controls (Fig. 2B,C). The activity of both TEP1 orthologs was similar to that of the isolated TIR domain of human SARM1, whereas the corresponding catalytic mutant of SARM1 (E642A) was inactive. No cyclic ADPR products were detected in reactions with TEP1 TeTIR under these conditions. To determine whether NADase activity is conserved among more divergent TEP1 homologs, we analyzed TeTIR domains from zebrafish (*Danio rerio*), the sponge *Amphimedon queenslandica*, and the amoeba *Dictyostelium discoideum*. All three wild-type proteins generated NAM and ADPR, whereas the corresponding glutamate mutants were inactive (Fig. 2D,E). Their activities were, however, lower than those observed for the human and mouse TEP1 TeTIR domains. In contrast, three prokaryotic TeTIR homologs tested under the same conditions did not show detectable activity (Suppl. Fig. S2B). Similarly, the isolated TeTIR domain of human NWD2 (residues 35–364) and mouse mNWD2 (residues 1-364) showed low activity, while human NPHP3 (residues 234-470) was completely inactive (Suppl. Fig. S2C).

**Figure 2:**
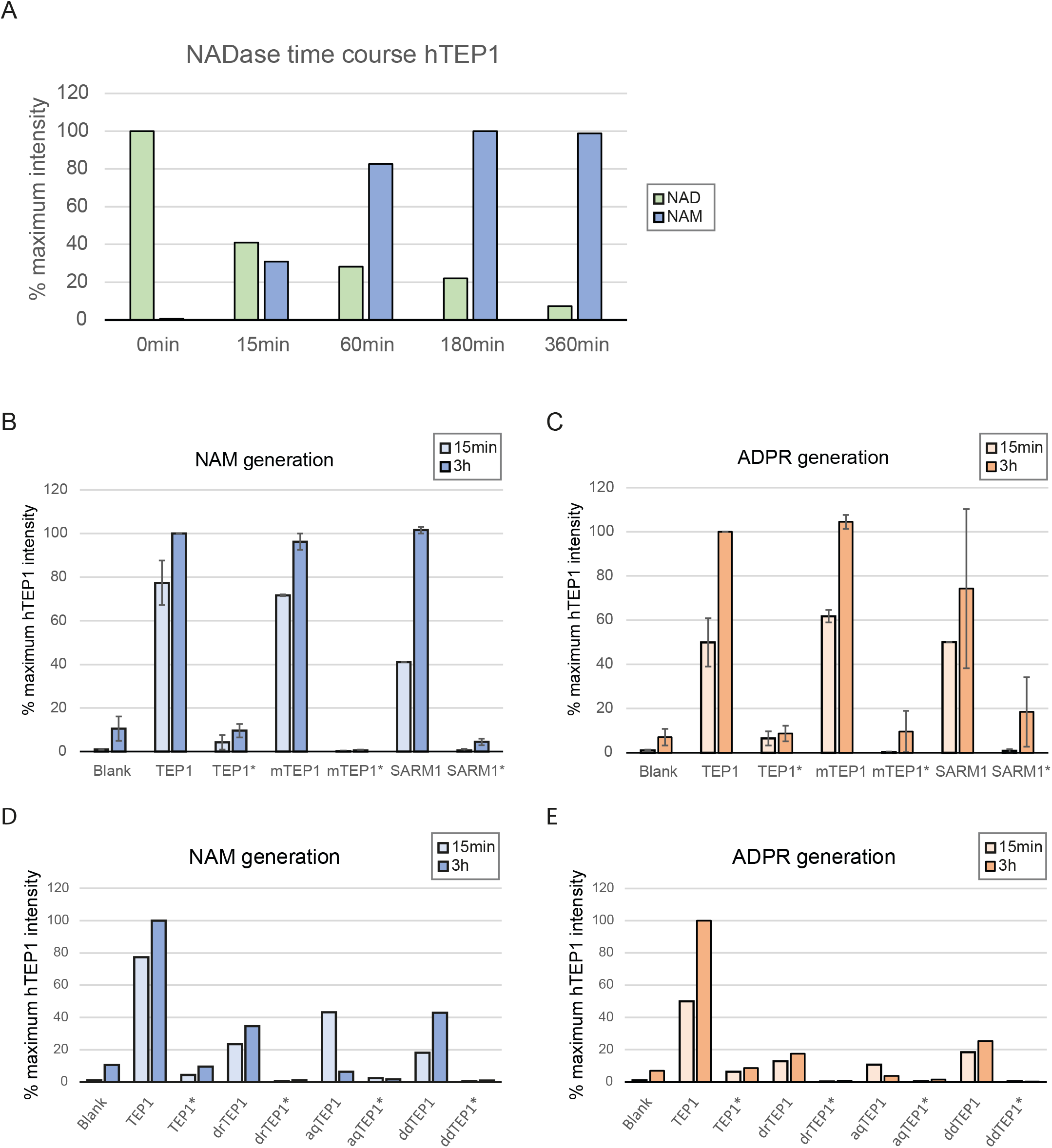
NADase activity of TeTIR domains. **(A)** Time course of NAD consumption (green) and NAM generation (blue) over 6 h. **(B, C)** NAM (blue) and ADPR (orange) generation, respectively, in NADase assays with human TEP1, murine mTEP1, and human SARM1, and their inactive glutamate mutants (TEP1*, mTEP1*, SARM1*). **(D, E)** NAM (blue) and ADPR (orange) generation, respectively, with human TEP1, *Danio rerio* drTEP1, *Amphimedon queenslandica* aqTEP1, *Dictyostelium discoideum* ddTEP1, and their corresponding inactive glutamate mutants (asterisks). For (B–E), measurements were performed after 15 min (light colors) or 3 h (dark colors).

Together, these data demonstrate that TeTIR domains from TEP1-like proteins possess NADase activity, which is conserved across diverse eukaryotic TEP1 homologs and dependent on the conserved catalytic glutamate. The activity of mammalian TEP1 TeTIR was comparable to that of SARM1-TIR, whereas substantially reduced or undetectable activity was observed in NWD2 and the prokaryotic TeTIR homologs.

### TEP1 TeTIR adopts an extended TIR-like fold

The biochemical characterization established that the TEP1 TeTIR domain possesses robust NADase activity comparable to that of SARM1-TIR. To investigate the structural basis of this activity and further assess the predicted relationship between TeTIR and TIR domains, we sought to determine the structure of the human TEP1 TeTIR domain. The wild-type domain did not yield suitable crystals, whereas the catalytically inactive glutamate mutant (TEP1*, E1008A) crystallized reproducibly. Crystallization was performed in the presence of NAD^+^ in an attempt to obtain a substrate-bound structure; however, the resulting crystals did not contain NAD^+^. The structure was determined at 1.3 Å resolution, with residues 895–1112 resolved except for residues 1083 and 1084. The asymmetric unit contained a single TeTIR molecule, and crystallographic and refinement statistics are summarized in Table 1. The crystal structure of TEP1* confirmed the predicted TIR-like fold, comprising a central parallel β-sheet surrounded by α-helices in a three-layer α/β/α sandwich (Figure 3A). Consistent with this assignment, a DALI search ^21^ identified significant structural similarity to multiple TIR domains as well as members of the MilB/DNPH family. Although SARM1-TIR was not among the top DALI matches, it yielded a significant Z-score and was selected for detailed comparison with TEP1-TeTIR because its NADase activity is well characterized.

**Table 1.**
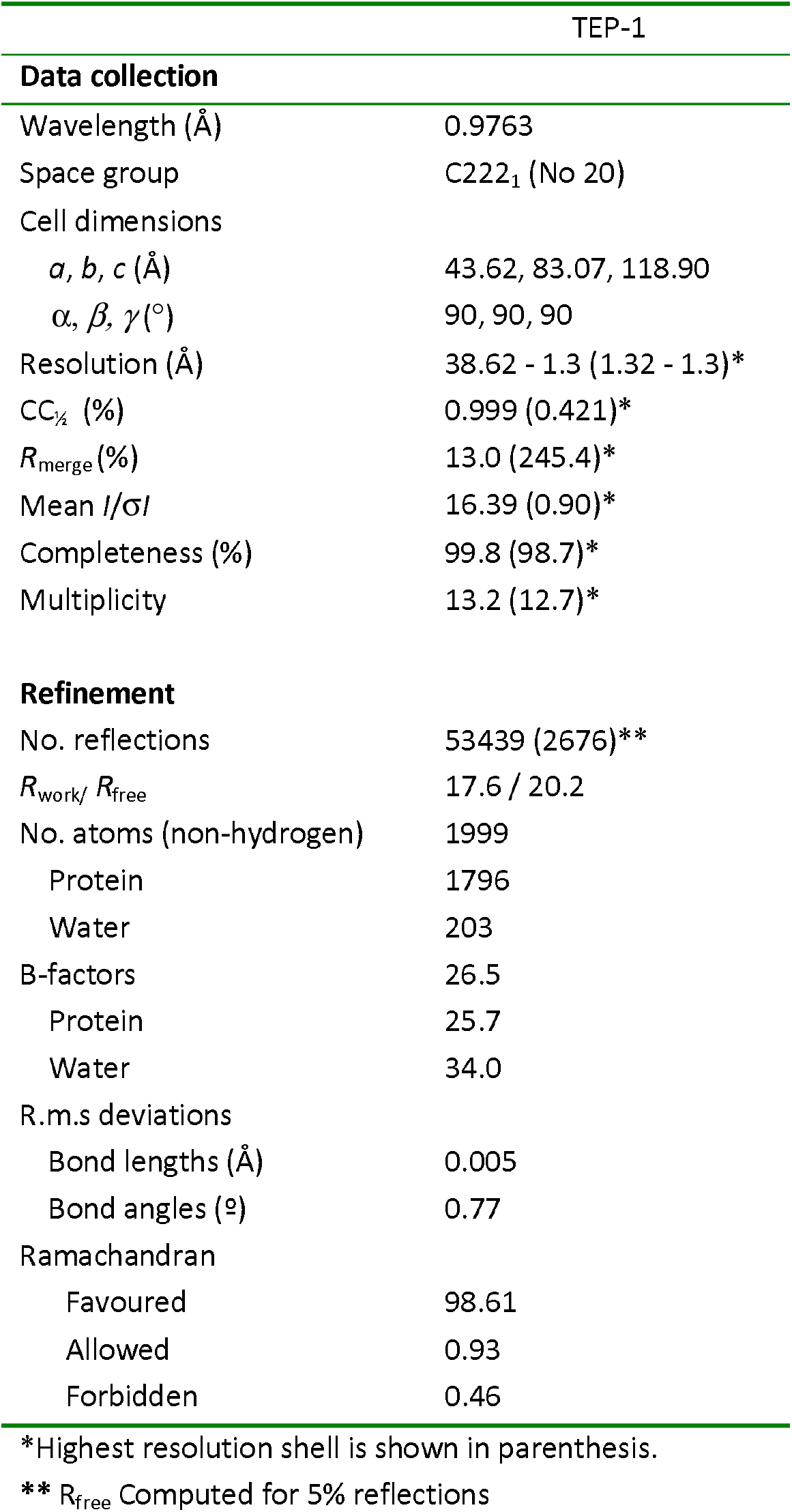
Data collection and refinement statistics.

**Figure 3:**
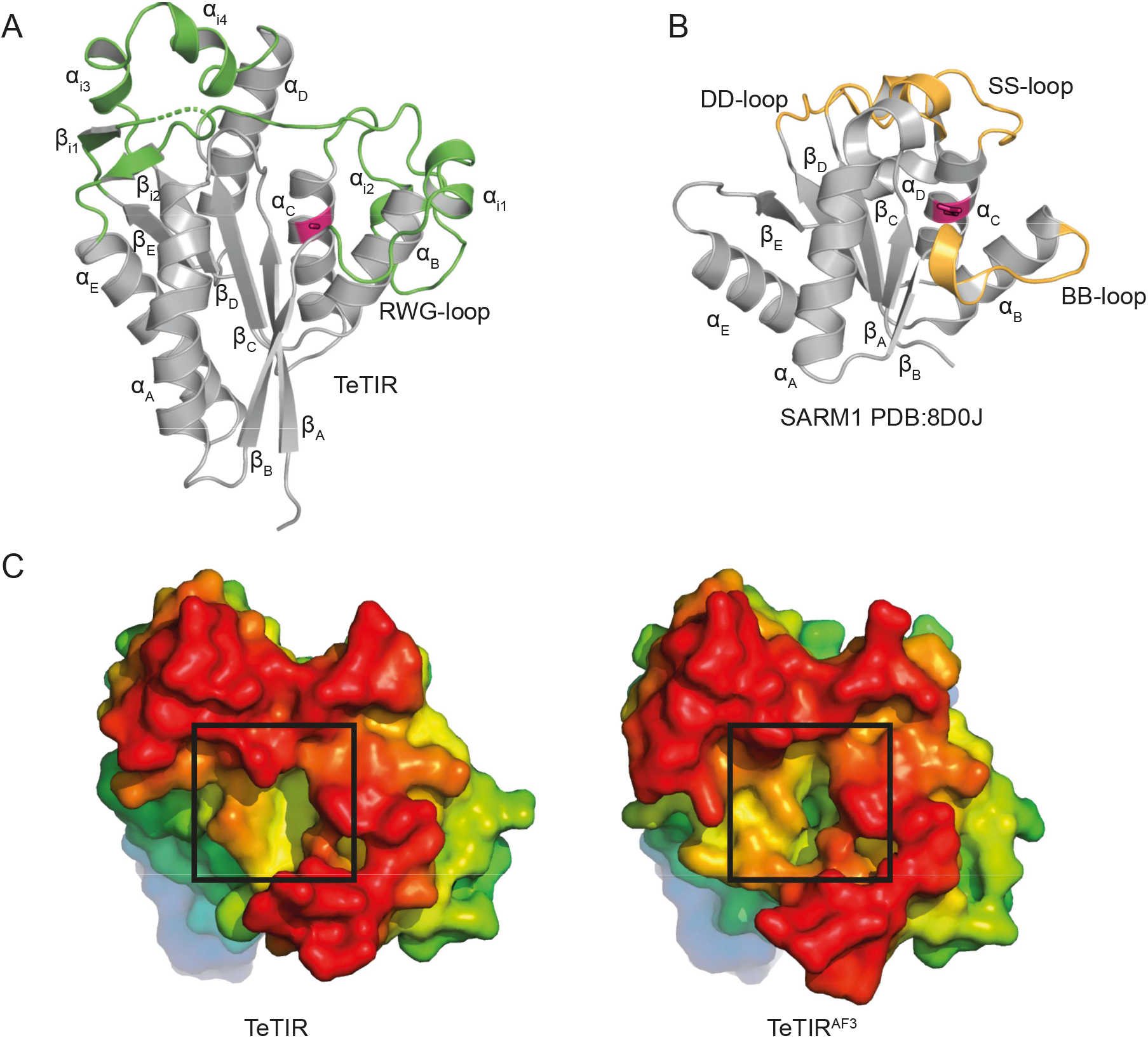
Crystal structure of TEP1 TeTIR. **(A)** Overview of the E1008A-mutant TeTIR structure (grey), with helices and strands labeled according to TIR nomenclature ^60^. Regions containing secondary-structure elements inserted into the TIR fold are shown in green. The two unresolved residues are indicated by a dotted line. Other deviations from the canonical TIR fold, such as extended helices and strands, are not highlighted. The catalytic glutamate, mutated to alanine in the structure, is shown in red. The flexible RWG-containing loop and inserted helix αi1 occupy the BB-loop region. **(B)** Apo structure of SARM1 (PDB: 8D0J) shown in the same orientation and using the same secondary-structure nomenclature. The catalytic glutamate is shown in red; loops implicated in SARM1 oligomerization and activation are shown in yellow. **(C)** Space-filling views of the E1008A-mutant TeTIR crystal structure (left) and the corresponding ligand-free AlphaFold 3 model (right). Surfaces are colored by depth. The difference in pocket openness is indicated by a black frame.

Superposition of the TEP1* and SARM1-TIR structures revealed a corresponding position of the catalytic residues, with SARM1 E642 occupying a position equivalent to TEP1 E1008 (Figure 3A,B). Despite the overall similar fold, the TeTIR structure contains several pronounced extensions, many of which correspond to additional secondary-structure elements inserted into the canonical TIR core (Figures 1B and 3A). The BB loop, which in SARM1 contributes to oligomerization-dependent activation of the TIR domain, is substantially extended in TeTIRs and adopts a distinct conformation. It also contains the highly conserved RWG motif, which is absent from canonical TIR domains. The CC loop, which in SARM1 contains the SARM1-specific (SS) region, is likewise substantially expanded in TeTIRs and contains an additional helix inserted before α_C_. The DD and EE loops are also markedly expanded, with two additional helices inserted before α_D_ and two additional β-strands inserted before α_E_, respectively. Whereas the core structural elements between β_A_ and α_D_ can be superimposed reasonably well between SARM1-TIR and TEP1-TeTIR, the C-terminal strand and helix adopt distinctly different arrangements. Sequence alignments and AlphaFold models indicate that these extensions and structural differences are conserved features of the TeTIR family rather than idiosyncratic features of TEP1. Together, these observations support the classification of TeTIRs as extended TIR-like domains.

Comparison of the crystal structure with the AlphaFold models revealed close agreement in the overall domain architecture, with the notable exception of the RWG-containing loop in the BB region (Supplementary Figure S3A). In the Alphafold models, E1008 projects into a pronounced cavity, whereas the corresponding region is largely occluded in the crystal structure of the E1008A mutant (Figure 3C).

### Structural basis of TeTIR active-site accessibility

The different conformations observed in the Alphafold model and crystal structure raise the question of how the experimentally observed NADase activity can be accommodated by the apparently closed conformation of the active-site region. In the Alphafold model, the catalytic residue E1008 is exposed within a pronounced pocket, whereas in the crystal structure of the catalytically inactive mutant, this region is largely occluded by the conformation of the RWG-containing loop and adjacent helix (Fig. 3C). We therefore investigated whether the two structures could represent alternative conformational states of the active-site region. Such a conformational transition might be induced by substrate binding or oligomerization. An oligomerization-dependent mechanism would be consistent with the position of the RWG-containing loop within the BB-region, which corresponds to the BB-loop of SARM1 and is implicated in the conformational regulation of SARM1 activity. However, purified wild-type TeTIR behaves as a monomer by size-exclusion chromatography, and the crystal structure is likewise monomeric, arguing against oligomerization being required for formation of the active conformation. Substrate-induced opening therefore remained a possible explanation, although NAD^+^ was already present during crystallization of the inactive E1008A mutant without producing an apparently open active-site conformation.

We therefore asked whether the catalytic glutamate itself might influence the conformational state of the active-site region. To address this question, we performed molecular dynamics (MD) simulations starting from the experimentally determined structure, either retaining the E1008A substitution or restoring E1008 in silico. To quantify changes in pocket accessibility, we monitored the volume of the pocket containing residue 1008 every 100 ps in the MD trajectories using Fpocket^22^. In simulations of the E1008A protein, the active-site region remained predominantly in the closed conformation (Fig 4A), with the RWG-containing loop preferentially adopting the conformation associated with pocket occlusion. In contrast, reintroduction of E1008 resulted in repeated and extended periods in which the pocket adopted an open conformation. Although the RWG-containing loop exhibited comparable conformational flexibility (Supplementary Fig S4A) in the two simulations, it preferentially sampled the open conformation in the presence of E1008. Consistent with the pocket-volume analysis, the relative solvent accessibility of residue 1008 showed the same E1008-dependent difference (Supplementary Fig. S4B). To further test whether E1008 influences this conformational equilibrium, we introduced the alanine substitution into an open-pocket conformation sampled from the E1008 trajectory. During subsequent MD simulation, the pocket progressively became more occluded, as reflected by a decrease in pocket volume and in the RSA of the catalytic residue (Fig. 4A, S4B). Together, these simulations suggest that E1008 shifts the conformational equilibrium of the active-site region toward the open, substrate-accessible state. This dynamic behavior provides a possible explanation for the absence of a defined ligand-binding pocket in the crystal structure and prompted us to investigate how NAD might be accommodated in the open conformation.

**Figure 4:**
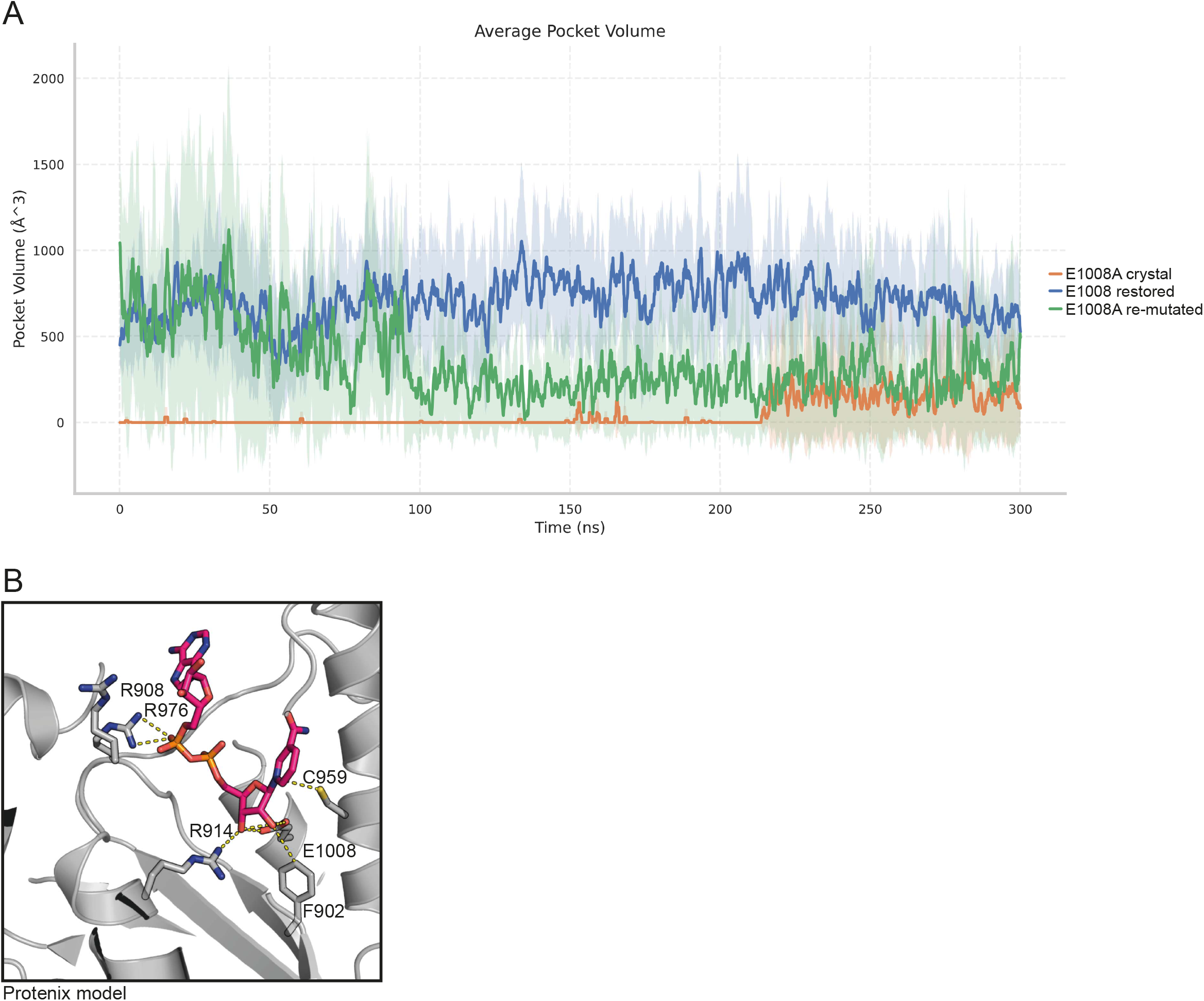
Structural basis of active-site accessibility. **(A)** Mean active-site pocket volume (i.e., the pocket containing residue 1008) over 300 ns MD trajectories. Each condition was simulated in three replicates: the original crystal structure (E1008A, orange), the crystal structure with E1008 restored in silico (blue), and an open-pocket structure sampled from the previous trajectory and subsequently re-mutated to E1008A (green). **(B)** NAD^+^ recognition in the Protenix model of TEP1-TeTIR. The protein is shown in grey cartoon representation and NAD^+^ as red sticks. Contact residues and the non-contacting R908 are shown as sticks.

Since our crystal structure does not contain NAD, we next sought to model how NAD could be accommodated in the active-site region and to identify residues that might contribute to substrate binding and catalysis. We first used AI-based structure prediction to model NAD-bound TEP1 TeTIR. The AF3 ^23^ and Protenix ^24^ models were broadly similar in their predicted NAD-binding configuration, but differed mainly in the RWG-containing loop (Supplementary Fig S4C), consistent with the conformational flexibility of this region observed in our MD simulations. The predicted NAD-bound models adopt a substrate-binding configuration resembling the ribose and base-recognition geometry observed in DNPH1 and SARM1-TIR structures (Fig. S4C and 1B). Several residues in this predicted binding site are conserved across the TeTIR family and occupy positions that have been implicated in catalysis or substrate recognition in these related enzymes. In particular, the catalytic glutamate E1008 is strictly conserved and occupies the position corresponding to the catalytic glutamates of DNPH1 and SARM1. The position of the second catalytic residue of DNPH1, D80, is instead occupied by the highly conserved C959 in TeTIR, which is positioned with its thiol group contacting the nitrogen of the NAM moiety. Finally, F902 is highly conserved in TeTIR and corresponds to the deoxyribose-binding Y24 of DNPH1; in the predicted models, F902 contacts the ribose of the NAM moiety, consistent with a possible role in substrate recognition.

The similarity of the predicted ribose–NAM binding mode to structures of DNPH1 and SARM1-TIR could potentially reflect the influence of structurally related protein–ligand complexes on AI-based structure prediction. We therefore sought an independent prediction of NAD binding using physics-based molecular docking. To account for conformational flexibility of the pocket, NAD was docked using an ensemble of structures sampled from a 1-µs MD simulation of the experimental structure with E1008 restored in silico. Representative conformations from four highly populated clusters were then used for NAD docking, and two visually distinct poses that retained the E1008–ribose contact were selected for each conformation and subjected to MD simulations. Unstable NAD-binding configurations were discarded, while configurations in which NAD remained associated with the protein were used as starting points for subsequent rounds of MD sampling and selection. This iterative procedure ultimately identified three configurations in which NAD remained stably bound throughout the MD trajectory, two of which adopted a binding mode similar to that predicted by AF3 and Protenix (Supplementary Fig. S4C). Analysis of these stable trajectories revealed recurring contacts between NAD and several TeTIR residues. In addition to E1008, C959 and F902, three conserved arginine residues, R914, R942, and R976, repeatedly contacted the NAD+ diphosphate and/or NAM-ribose, suggesting a role in NAD recognition. In contrast, R908, which occupies a position similar to residues involved in phosphate recognition in SARM1-TIR, is poorly conserved across the TeTIR family and showed fewer contacts with the NAD+ diphosphate in our models.

### Mutagenesis identifies residues required for TeTIR NADase activity

Together, these analyses identified E1008, C959, F902, R914, R942, and R976 as candidate residues involved in catalysis or substrate recognition. We also considered several additional residues based on their conservation and structural position. W943 is a neighbor of R942 in the conformationally flexible RWG loop and is unusually well conserved despite not forming apparent contacts with either the substrate or the remainder of the TeTIR domain. Instead, W943 points outward into solvent-exposed space in the experimental structure and in most of the predicted models. This unusual exposure suggested that W943 might participate in an intermolecular interface, potentially contributing to TeTIR oligomerization. We therefore included W943 as a candidate residue that might influence the conformation of the RWG-containing loop and the associated open–closed transition of the active-site region, while also testing the possibility that they contribute to intermolecular interactions. Finally, R908 was selected because it occupies a position analogous to residues involved in phosphate recognition in SARM1-TIR and resides in a loop adjacent to the predicted NAD-binding site. However, R908 does not appear to contact the NAD phosphates in most models and is less conserved than R914 and R976. We therefore predicted that R908 would contribute less directly to substrate recognition than the two highly conserved arginine residues R914 and R976.

As shown in Fig. 5, mutations of the proposed catalytic or substrate-binding residues resulted in a drastic reduction in activity, approaching the background level observed for the inactive E1008A mutant. In particular, the near-complete loss of activity upon mutation of C959 and F902 supports their model-derived assignment as a catalytic residue and a determinant of ribose recognition, respectively. R914A and R976A were similarly inactive, consistent with their predicted roles in contacting the NAM-proximal ribose or the NAD phosphates. In contrast, R908A retained considerable residual activity despite a reduction relative to the wild-type protein, suggesting that R908 is less critical for NAD binding than R914 and R976. Among the residues of the RWG-containing loop, R942A was inactive, whereas W943A caused only a modest reduction in activity. Finally, the S905E mutant designed to be sterically disruptive was also devoid of activity. Because S905 lines the predicted substrate-binding pocket, introduction of the larger glutamate side chain was expected to interfere with substrate accommodation; accordingly, this result was interpreted as evidence for a steric effect rather than as evidence for a direct catalytic or substrate-recognition role of S905.

**Figure 5:**
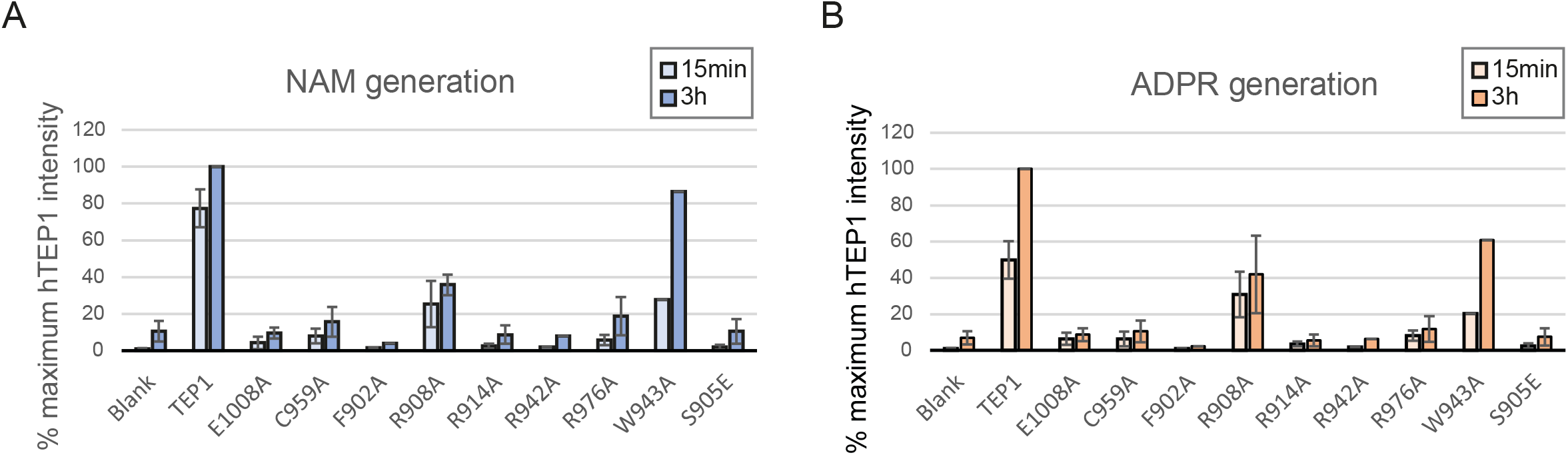
Functional analysis of residues involved in NAD+ recognition and catalysis. **(A, B)** NAM (blue) and ADPR (orange) generation, respectively, in NADase assays with wild-type human TEP1 and the indicated mutants. The blank represents an enzyme-free assay. Measurements were performed after 15 min (light colors) or 3 h (dark colors).

### TEP1 NADase activity leads to NAD depletion in cells

SARM1 and several bacterial TIR proteins have been shown to cause pronounced NAD^+^ depletion that can lead to cellular toxicity. For SARM1, NADase activity is enhanced by oligomerization, including through the SAM domains, but the isolated TIR domain can also cause NAD^+^ depletion and cell death when expressed in cells. Bacterial TIR proteins likewise possess intrinsic NADase activity, and some have been shown to deplete NAD^+^ in cells. We therefore asked whether isolated TeTIR domains are sufficient to deplete cellular NAD^+^. To this end, we expressed the TeTIR domains of human TEP1 and homologs from several other species in HEK293 cells and measured cellular NAD+ levels 24 h after transfection. As shown in Fig. 6A,B, the TeTIR domains of human and mouse TEP1 reduced cellular NAD^+^ levels to approximately 30–40% of those in control transfections. This effect depended on the catalytic glutamate. Expression of human SARM1 resulted in a similar reduction in cellular NAD^+^ levels, whereas the corresponding glutamate mutant was ineffective (Fig. 6A). The bacterial TIR domain of TcpC likewise reduced cellular NAD^+^ levels to approximately 40% of control levels upon overexpression, whereas the TIR domain of the plant protein RPP1 did not measurably deplete cellular NAD^+^ (Fig. 6B). Several mutated versions of hTEP1 that showed strongly reduced NADase activity in vitro were also tested and likewise failed to cause NAD^+^ depletion below control levels (Supplementary Fig. S5A).

**Figure 6:**
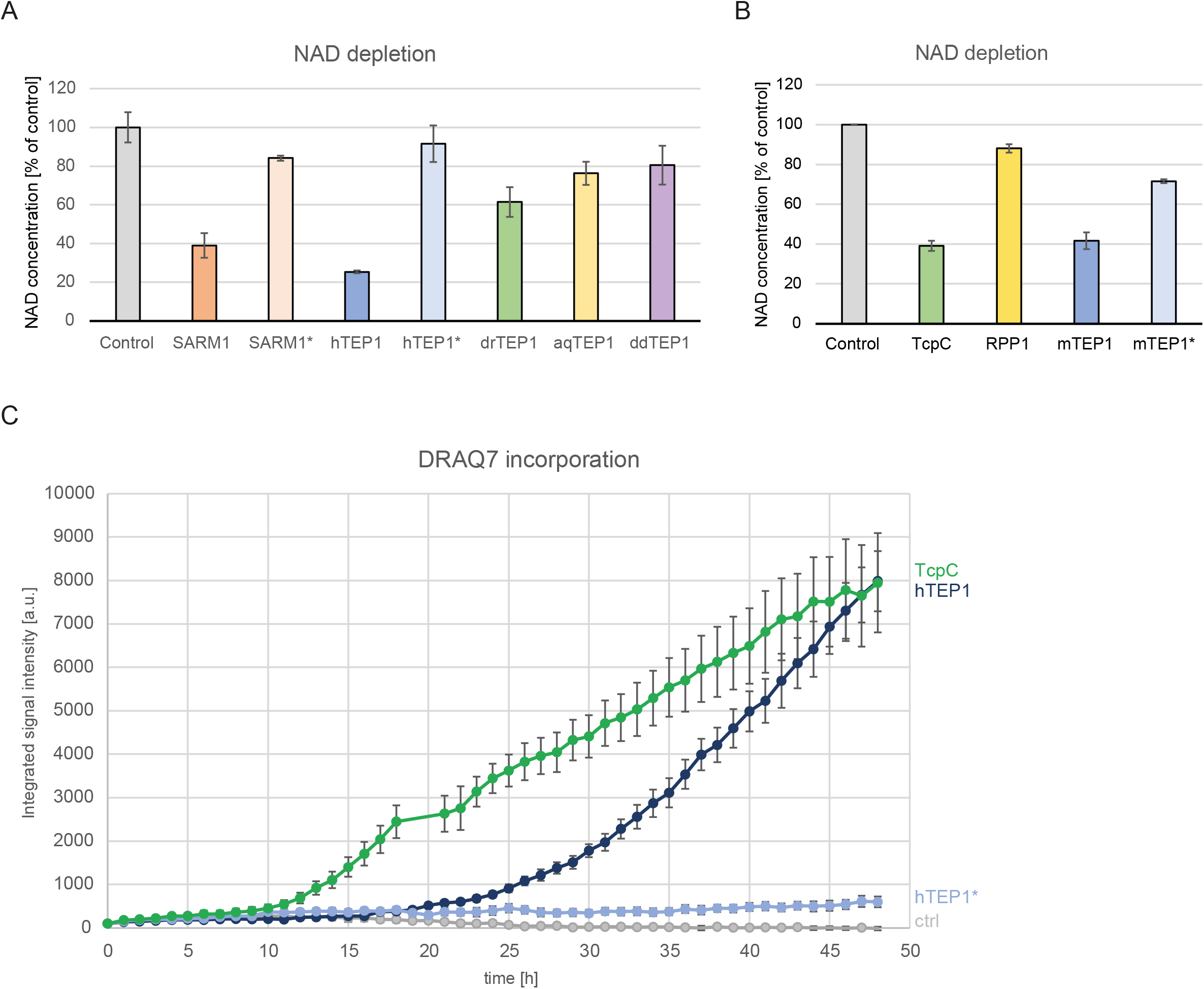
Cellular NAD depletion and cell death induced by TeTIR overexpression. **(A, B)** Cellular NAD(H) content measured 24 h after transfection with the indicated constructs. Untransfected cells served as controls. **(A)** Human SARM1 and TEP1 and their inactive glutamate mutants, together with drTEP1, aqTEP1, and ddTEP1. **(B)** mTEP1 and its inactive glutamate mutant, and the TIR domains of bacterial TcpC and plant RPP1. **(C)** Time course of DRAQ7 incorporation over 48 h following transfection with untransfected control (grey), inactive hTEP1* (light blue), active hTEP1 (dark blue), or bacterial TcpC (green), measured by Incucyte imaging. The integrated signal intensity is shown.

During these experiments, we noticed that cells overexpressing enzymatically active TeTIR domains showed increased cell death compared with cells expressing inactive variants. To investigate this phenotype more closely, we performed live-cell imaging of HEK293 cells expressing various TeTIR constructs and monitored cell death by incorporation of DRAQ7, a nuclear dye that enters cells upon loss of membrane integrity. As shown in Fig. 6C, expression of the wild-type human TEP1 domain resulted in substantial DRAQ7 incorporation beginning approximately 20 h after transfection and increasing throughout the 48 h observation period. In contrast, the inactive E1008A mutant (TEP1*) did not induce detectable cell death. Expression of the NAD-depleting TIR domain of bacterial TcpC caused a similar extent of cell death, although with an earlier onset beginning at approximately 10 h after transfection (Fig. 6C). Among the other TeTIR domains tested, mouse TEP1 was even more effective than human TEP1. In contrast, the TeTIR domains of human and mouse NWD2 caused only marginal cell death, whereas the enzymatically inactive TeTIR domains of TTC41 and NPHP3 did not show a detectable toxic effect (Supplementary Fig. S5B).

## Discussion

Our identification of TeTIR domains expands the known repertoire of TIR-like NADases and reveals a previously unrecognized occurrence of this enzymatic activity in STAND-type proteins. TeTIR domains are related to TIR domains at the sequence and structural levels and retain the conserved catalytic glutamate characteristic of TIR-like NADases, while being substantially more divergent and structurally elaborated than canonical TIR domains. In TEP1 and related proteins, the TeTIR domain is located immediately upstream of the central NACHT ATPase, thereby occupying the position of an effector domain within a STAND-protein architecture. This organization is shared by the mammalian TEP1, NWD1, NWD2, TTC41, and NPHP3 proteins and is also found in related proteins from other eukaryotes and bacteria. TEP1 is additionally characterized by an N-terminal TROVE/VWA region preceding the TeTIR domain. The coupling of an effector domain to a central STAND ATPase is a recurring feature of signaling proteins that assemble into higher-order complexes, including inflammasomes, apoptosomes, and plant resistosomes, in which nucleotide-dependent activation of the STAND domain is linked to activation or assembly of the associated effector machinery. Although no such activation mechanism or higher-order assembly has yet been demonstrated for TEP1 or other TeTIR-containing proteins, the conserved placement of TeTIR immediately adjacent to the NACHT domain suggests that these proteins may likewise couple nucleotide-dependent conformation changes to control of their NADase effector domain. The induction of TEP1 expression by interferon ^25^ further places this protein in an inducible immune-response context, consistent with a sensor-effector role.

The catalytic activity of TeTIR domains is not uniformly conserved across the family. While TEP1 homologs from several eukaryotic species showed detectable NADase activity, NWD2 displayed only weak activity, and other TeTIRs including the prokaryotic homologs were inactive under the conditions used. These differences could reflect genuine divergence in catalytic function, but an alternative possibility is that activity of some TeTIR domains depends on structural or regulatory features absent from the isolated domains used in our assays. In particular, given their association with NACHT-containing STAND proteins, activation-dependent oligomerization or interactions with other regions of the full-length proteins could potentially contribute to formation or stabilization of an active TeTIR conformation. TEP1 and related proteins containing N-terminal TROVE domains might also be regulated by RNA binding, as this region has been shown to bind RNAs with limited sequence specificity, including vault and telomerase RNAs ^26^.

The structural similarity of TeTIR to both TIR NADases and the MilB/DNPH family suggests that these proteins may share features of substrate recognition and cleavage chemistry, with the catalytic glutamate residue (E1008 in human TEP1) being a conserved key feature. Although SARM1 and DNPH1 cleave analogous N-glycosidic bonds in related substrates, different catalytic mechanisms have been proposed. DNPH1 was reported to employ a two-step “double-displacement” mechanism, initiated by D80 protonating the nitrogen of the leaving group and followed by nucleophilic attack of the catalytic glutamate on the ribose carbon formerly bound to the protonated nitrogen. In the final step, D80 activates a water molecule for hydrolysis of the resulting glycosyl-enzyme intermediate ^19^. By contrast, a “ternary-complex” mechanism has been proposed for the TIR domain of SARM1, in which water (or another nucleophile) attacks the NAM-bound ribose carbon, assisted by hydrogen bonds between the catalytic glutamate and the two hydroxyl groups of the NAM-proximal ribose ^27^. Based on residue conservation and their predicted positions, TeTIRs appear compatible with a SARM1-like one-step mechanism but lack the protonating aspartate of DNPH1. Whereas SARM1 and other classical TIR domains completely lack a functional residue at this position, TEP1 and its orthologs contain a conserved cysteine. Although cysteine would not be expected to protonate the leaving-group nitrogen in the manner proposed for DNPH1 D80, C959 is positioned close to this atom in our substrate-bound models, and its replacement by alanine abrogates enzymatic activity. Thus, C959 may participate in another step of catalysis or contribute to substrate binding or positioning, although its precise role remains unclear.

The phosphate-binding region provides a further distinction between TeTIR and SARM1. In the SARM1 TIR domain, R569 and R570 contribute to recognition of the diphosphate moiety of NAD+. In TeTIR, no arginine pair occupies the corresponding position. R908 is located near this region but is somewhat displaced relative to the SARM1 residues. Although R908 does not contact the phosphates in our structural models, some of the MD simulations showed substantial phosphate interactions. However, R908 is not well conserved among TeTIRs, and its mutation only partially reduced activity, arguing against a central role in phosphate recognition. In contrast, R976 emerges as a prominent candidate for direct phosphate recognition. It is predicted to contact the NAM-distal phosphate in all of our substrate-bound models, and these interactions are also maintained throughout the MD simulations. Consistent with an important functional role, R976A completely abolished NADase activity. Another conserved arginine, R914, provides a different case: although it does not generally contact the phosphates, it is positioned near the NAM-linked ribose in most models and is frequently found in this vicinity during the MD simulations. Its strong mutational phenotype therefore suggests an important role in NAD+ binding or positioning rather than direct phosphate recognition. Finally, R942, located within the RWG loop, does not contact NAD+ in the static models but frequently approaches the NAM-proximal phosphate during the MD simulations. The loss of activity upon R942A substitution suggests that this otherwise structurally mobile region contributes to formation or stabilization of the NAD+-binding site. Together, these observations indicate that TeTIRs retain the general active-site architecture of TIR-like NADases while using a distinct arrangement of residues for NAD+ recognition and positioning, including contributions from the structurally mobile RWG loop.

An additional feature emerging from the structural and mutational analyses is the apparent coupling between the catalytic glutamate and active-site accessibility. Rather than functioning solely as the catalytic residue, E1008 appears to influence the conformational state of the surrounding active-site region. This may provide a mechanism by which formation of a productive substrate-binding site is coupled to the catalytic machinery. Such conformational plasticity could also provide a potential point of regulation in the context of the full-length STAND protein. Although isolated TEP1 TeTIR is active without detectable oligomerization, interactions with the NACHT domain, the N-terminal TROVE/VWA region, or other cellular factors could regulate the equilibrium between substrate-accessible and occluded conformations. If oligomerization contributes to activation of full-length TeTIR-containing STAND proteins, it is unlikely to involve the same interface characterized in SARM1, as the corresponding surface regions are poorly conserved between SARM1 and TeTIRs. Nevertheless, the structurally dynamic RWG loop occupies a position close to the BB loop of SARM1, which has an important role in the assembly and activation of SARM1 TIR domains. Together with the functional importance of R942 within this loop, this positional correspondence raises the possibility that the RWG region could contribute to intermolecular interactions or conformational changes associated with TeTIR activation. Whether this region participates in oligomerization or instead primarily contributes to active-site organization remains to be determined.

The biochemical activity of TeTIRs also has clear consequences when the isolated domains are expressed in cells. Expression of the human and mouse TEP1 TeTIR domains caused a pronounced reduction in cellular NAD^+^ levels, comparable to that observed upon expression of the SARM1 TIR domain, whereas catalytically inactive mutants did not deplete NAD^+^. These findings demonstrate that TeTIR NADase activity is sufficient to perturb cellular NAD^+^ homeostasis when the catalytic domain is strongly expressed. However, these experiments do not establish that NAD^+^ depletion is a physiological function or mechanism of TEP1. The isolated TeTIR domain is expressed independently of the regulatory context provided by the full-length STAND protein, and the magnitude and timing of overexpression may substantially exceed those encountered under endogenous conditions. Thus, the cellular experiments primarily demonstrate the capacity of TeTIR activity to consume cellular NAD^+^ and, at sufficiently high activity, to produce a toxic phenotype.

The reduction in cellular NAD^+^ levels was accompanied by increased cell death upon expression of active TEP1 TeTIR, whereas catalytically inactive variants showed little or no such effect. This correlation is consistent with NAD^+^ depletion contributing to the toxic phenotype, but does not establish the mechanism by which TeTIR activity compromises cell viability. The cellular phenotype therefore demonstrates that TeTIR NADase activity can be cytotoxic when strongly expressed. The delayed onset of cell death may in part reflect the dynamic balance between NAD^+^ consumption by the expressed NADase and ongoing cellular NAD+ biosynthesis. Cellular NAD^+^ pools are continuously replenished, and this capacity could initially compensate for TeTIR-mediated NAD^+^ hydrolysis. With sustained NADase activity, however, consumption may eventually exceed the rate of replenishment, leading to progressive NAD^+^ depletion and, ultimately, loss of cell viability. This provides a possible explanation for the lag between TeTIR expression and the onset of cell death, although the contribution of NAD^+^ depletion to the timing of the phenotype remains to be established.

The localization of TEP1 within the vault complex also raises an intriguing possibility concerning its relationship to PARP4, the other enzymatically active minor vault component ^28, 29^. PARP4 is an NAD^+^-consuming ADP-ribosyltransferase, and recent work has implicated functional interplay between SARM1 and PARP1 in parthanatos, illustrating how distinct NAD-consuming enzymes can participate in a common cellular response ^30^. Whether TEP1 and PARP4 are similarly functionally coupled is unknown, but their coexistence within the vault raises the possibility that TEP1 NADase activity could influence local NAD^+^ or ADP-ribose metabolism and thereby modulate PARP4 activity. This remains entirely speculative, however, and determining whether TEP1 NADase activity is functionally linked to PARP4 or other vault functions will require direct investigation.

## Materials and Methods

### Sequence analysis & structure prediction

Sequence alignments within protein families were generated using the L-INS-I method implemented in the MAFFT package ^31^. Sequence alignment between the distantly related TIR/DNPH/TeTIR families were generated by superposition of their structures using DALI ^21^. Generalized profiles were derived from multiple alignments and searched against the UniProt database ^32^ using pfsearchV3 ^33^. HMM-to-HMM searches to establish sequence similarity between protein families were carried out using the HHSEARCH method ^17^. The initial structure predictions of isolated proteins and domains were performed using the Alphafold-2 server ^20^. Predictions of multi-protein assemblies and protein-ligand complexes were performed using a local installation of Alphafold-3 ^23^ with five seeds per run. Additional protein-ligand complexes were predicted using the Protenix server ^24^. Energy minimization was performed using YASARA v25.1.13 with the YASARA force field ^34^.

### Structure Editing and molecular docking

The catalytic glutamate was computationally restored in the experimental E1008A structure using the PyMOL mutagenesis wizard, replacing A1008 with glutamate (A1008E). Conversely, an open E1008 conformation sampled from the molecular dynamics trajectory was computationally mutated from E1008 to alanine (E1008A) using the PyMOL mutagenesis wizard. Formal charges, bond definitions, and missing hydrogens in the docked NAD+ poses were added using Discovery Studio Visualizer. PDBFixer ^35^ was used to process all starting protein configurations by removing heteroatoms and rebuilding missing heavy atoms.

Molecular docking was carried out using AutoDock Vina through the AMDock GUI (v1.5.2) at an exhaustiveness of 62. The search space was limited by manually selecting a 27 × 21 × 27 Å box centered on the catalytic glutamate. The receptor and ligand were protonated at pH 7.0 using AutoDock Tools prior to docking. Docked poses were manually inspected for consistency with known structure–activity relationships (SARs), and two visually distinct poses per cluster were selected.

### Molecular Dynamics Simulations

Molecular dynamics (MD) simulations were carried out using the GROMACS 2024.1 engine ^36^ with the CHARMM36 force field ^37^. NAD^+^ parameters were generated using the CGenFF engine ^38^. Prior to simulation, the protonation states of all titratable amino acids in the starting protein structures were assigned using PROPKA 3 ^39^ at pH 7.0. The starting configurations were placed in a dodecahedral box containing CHARMM-modified TIP3P water ^40^, and the appropriate number of counterions was added to neutralize the system. The systems were energy-minimized using the steepest-descent algorithm with a maximum force threshold of 200 kJ mol^−1^ nm^−1^, followed by 100 ps NVT equilibration using a V-rescale thermostat ^41^ and a 5 ns fully restrained NPT equilibration using a C-rescale barostat ^42^. Unrestrained production simulations were carried out in the NPT ensemble using a V-rescale thermostat and Parrinello–Rahman barostat. All simulations were performed at a target temperature of 300 K and a target pressure of 1 bar, using LINCS constraints ^43^ and a 2-fs timestep. Particle Mesh Ewald ^44^ was used for long-range electrostatic interactions, while van der Waals interactions were treated using a cutoff scheme with a force-switch modifier. SETTLE constraints algorithm was used for treatment of rigid water molecules ^45^. Independent replicates were generated by assigning new random velocities prior to each production simulation.

Each combination of sampled receptor geometry and docked pose was subjected to 100 ns of MD simulation in five independent replicates. Replicates meeting the survival criteria were subsequently extended by initiating 500 ns simulations from their respective endpoints, using three independent replicates for each selected endpoint. The same selection and extension procedure was applied in a third round, using 500 ns simulations in three independent replicates. The endpoints of the resulting trajectories were then analyzed, and distinct models were subjected to 300 ns simulations in five independent replicates to generate trajectories for PLIF quantification.

### Analysis of MD Simulations Results

All trajectories were first processed to remove artifacts arising from periodic boundary conditions and aligned to the starting structure prior to analysis. For trajectories of protein–ligand complexes, the interatomic distance between the OE1 atom of Glu1008 and the H51 atom of the NAM-side ribose was calculated for all frames. This distance was used as a stability metric, with distances below 4.5 Å heuristically classified as indicating a bound state and distances above 4.5 Å as indicating an unbound state. RMSD-based clustering using the GROMOS method of Daura et al. ^46^ with a 1.7 Å cutoff was performed on the heavy atoms of residues lining the putative binding site to identify representative receptor conformations. Backbone RMSD was calculated using the starting structure as the reference. Per-residue backbone RMSF profiles were calculated over the full trajectory after fitting to the first frame.

Protein–ligand interaction fingerprints (PLIFs) were calculated using the ProLIF ^47^, MDAnalysis ^48^, and RDKit (https://www.rdkit.org) Python libraries for every 10th frame of each trajectory, corresponding to a 100 ps sampling interval. PLIFs were quantified across replicates by calculating the frequency of occurrence of each interaction as the number of frames in which the interaction was detected divided by the total number of analyzed frames.

Accessibility of the putative binding site was evaluated by calculating relative solvent accessibility (RSA) and using the fpocket cavity detection algorithm ^22^. Every 10th frame of the relevant trajectories (100 ps interval) was extracted as a separate PDB file. The Python interface of the open-source version of PyMOL ^49^ was used to calculate the RSA of the catalytic residue Glu1008. The fpocket package was used with default settings to detect pockets in each frame. The atomic composition of the fpocket-predicted cavities was analyzed using Python, and the volume of pockets containing the catalytic residue Glu1008 was extracted. Frames in which no viable pocket containing Glu1008 was detected were assigned a pocket volume of 0.0.

### Cloning & mutagenesis

All coding regions of TeTIR protein were obtained by gene synthesis (IDT) and cloned into the pOPIN-S ^50^and pcDNA5 vector using the In-Fusion HD Cloning Kit (Takara Clontech). All constructs from non-mammalian sources have been codon optimized. Point mutations were introduced using the QuikChange Lightning kit (Agilent Technologies). A list of all primers and constructs used are provided in Supplementary Table 1.

### Protein expression and purification

All proteins were expressed from the pOPIN-S vector with an N-terminal 6His-SMT3 tag ^50^. Constructs encoding TEP1 residues 895–1117, SARM1 residues 559–700, and the point mutants other than the catalytic glutamate mutants were transformed into *Escherichia coli* BL21-AI (Invitrogen). Cultures of 6–12 L were grown in LB medium at 37 °C to an OD_600_ of 0.8, cooled to 18 °C, and induced with 1 mM isopropyl β-D-1-thiogalactopyranoside (IPTG) and 20 mL of a 20% arabinose solution. Constructs encoding hTEP1-E1008A (hTEP1*), mTEP1-E1017A (mTEP1*), and SARM1-E642A (SARM*) were transformed into *E. coli* Rosetta (DE3) pLysS. 6-L cultures were grown in LB medium at 37 °C to an OD_600_ of 0.8, cooled to 18 °C, and induced with 0.1 mM IPTG.

After 16 h, cells were harvested by centrifugation at 5,000 × *g* for 15 min. Following freeze-thawing, cell pellets were resuspended in binding buffer (300 mM NaCl, 20 mM Tris pH 7.5, 20 mM imidazole, and 2 mM β-mercaptoethanol) containing DNase and lysozyme and lysed by sonication using 10-second pulses at 50 W for a total sonication time of 6 min. Lysates were clarified by centrifugation at 50,000 × *g* for 1 h at 4 °C, and the supernatants were subjected to affinity purification on HisTrap FF columns (Cytiva) according to the manufacturer’s instructions. The 6His-SMT3 tags were removed by incubation with SENP1(415–644). Proteins were then dialyzed against binding buffer, and the cleaved affinity tags together with His-tagged SENP1 were removed by a second round of affinity purification on HisTrap FF columns. All proteins were subjected to final size-exclusion chromatography on a HiLoad 16/600 Superdex 75 pg column equilibrated in 20 mM Tris pH 7.5, 150 mM NaCl, and 2 mM dithiothreitol (DTT). Proteins were concentrated using Vivaspin 20 centrifugal concentrators (Sartorius), flash-frozen in liquid nitrogen, and stored at −80 °C. Protein concentrations were determined from A_280_ measurements using theoretical extinction coefficients calculated from the amino acid sequences.

### In vitro NADase assays by LC-MS

Purified recombinant protein (10 μg) was incubated with 100 μM NAD^+^, 1 mM MgSO4 in a total volume of 100 μl of SEC buffer for 15 min or 3h at 37°C. Reactions were stopped by freezing the sample at −20 °C. 20 µl methanol was added to the sample before centrifugation at 4 °C for 10 min at full speed. Chromatography was performed on a Nexera XR 40 series HPLC (Shimadzu) using a Nucleodur Sphinx RP, 3 µm, 150 x 2 mm (Macherey Nagel). Samples (4 μl) were injected at a flow rate of 0.3 ml/min using 0.1% formic acid and acetonitrile as mobile phases A and B, respectively. Metabolites were eluted using the 20 min gradient profile 0 min, 1% B; 0-5 min, 100% B; 5-8 min 100% B, 8-8.1 min, 1% B. The LCMS-8060 triple quadrupole mass spectrometer with electro spray ionization (Shimadzu) was operated in positive mode. Scheduled multiple reaction monitoring (MRM) was used to monitor analyte parent ion to product ion formation. MRM conditions were optimized using authentic standard chemicals including: NAD^+^([M+H] 664.00>136.00, 664.00>427.90, 664.00>523.95), ADPR ([M+H] 559.80>136.35, 559.80>348.10, 559.80>97.20), nicotinamide ([M+H] 123.00>80.00, 123.00>78.00, 123.00>53.00), cADPR ([M+H] 541.80>136.00, 541.80>428.10, 541.80>348.15), 2′cADPR or 3′cADPR ([M+H] 541.90>136.00, 541.90>427.95, 541.90>97.05). Both Q1 and Q3 quadrupoles were maintained in unit resolution. LabSolutions LC-MS v5.118 software was used for data acquisition and LabSolutions Postrun for processing (both Shimadzu). Metabolites were quantified by scheduled MRM peak integration. Figures were done on excel.

### Cell culture and Transfection

HEK293T cells obtained from ATCC were cultured at 37 °C and 5% CO2 in Dulbecco’s modified Eagle’s medium (DMEM; Gibco), supplemented with 10% fetal bovine serum (FBS) and 1% penicillin-streptomycin. Cells were grown to 90% confluence before harvesting. Cell pellets were resuspended in lysis buffer (20 mM Tris pH 7.5, 150 mM NaCl, 0.5% NP-40, 2 mM EDTA, and 5 mM N-ethylmaleimide [NEM]) and sonicated using three 10-second pulses. Cell debris was removed by centrifugation at 16,000 × *g* for 15 min at 4 °C. Unreacted NEM was quenched by addition of 10 mM DTT.

Plasmid DNA was incubated in serum-free DMEM for 10 min at room temperature. Polyethylenimine (PEI) transfection reagent was added and the mixture was incubated for a further 10 min. Before transfection, the cell culture medium (complete DMEM) was replaced with DMEM containing 5% FBS. The transfection mixture was added to the cells for 4–5 h, after which it was replaced with fresh complete DMEM.

### Western Blots

The cell pellets were resuspended in 2× or 5× Laemmli buffer and boiled at 95 °C for 5 min. Samples were separated on 12% Tris-glycine gels and transferred to PVDF membranes by semi-dry Western blotting. Membranes were probed with either anti-HA primary antibody (Miltenyi Biotec, 130-091-972; 1:2000) overnight at 4 °C or for 1 h at room temperature, or with HRP-conjugated anti-DYKDDDDK antibody (Miltenyi Biotec, 130-101-572; 1:4000) overnight at 4 °C or for 1 h at room temperature, in 5% milk in PBS-T. For detection of HA-tagged proteins, membranes were subsequently incubated with HRP-linked anti-mouse secondary antibody (Cell Signaling Technology, Cat. #7076; 1:3000) in 5% milk in PBS-T for 1 h at room temperature. HRP activity was detected using WesternBright chemiluminescent reagent (Advansta, K-12045). Blots were imaged using Image Lab software version 5.2.1.

### Cellular NAD assays

NAD(H) concentrations in transfected cells were determined using the NAD^+^/NADH Assay Kit (Sigma-Aldrich). Cells were harvested 24h after transfection, as described above. Protein concentrations were determined using a Bradford assay (Roti-Quant; Roth) and used for normalization. Cell pellets were then heated at 60 °C for 5 min, followed by addition of 20 µL Assay Buffer and 100 µL NADH Extraction Buffer. Samples were briefly vortexed and centrifuged at 14,000 × g for 5 min. The supernatants were used for NAD^+^/NADH quantification. For the assay, 50 µL of each sample or NAD^+^ standard was transferred to a black, flat-bottom 96-well plate. The Working Reagent was freshly prepared immediately before the assay and contained a lactate dehydrogenase cycling reaction, in which the formed NADH reduces a probe to generate a fluorescent product. Fifty µL of Working Reagent was rapidly added to each well. The plate was gently mixed, and fluorescence intensity was immediately measured at an excitation wavelength of 530 nm and an emission wavelength of 585 nm. The plate was then incubated at room temperature for 10 min protected from light, followed by a second fluorescence measurement. Measurements were performed using a TECAN plate reader in kinetic mode with shaking and a manually adjusted gain of 44.

### Cell death assays

Cells were seeded at 30,000 cells per well in 48-well plates. The following day, cells were transfected with the respective constructs using PEI. After 5–6 h of incubation, the culture medium was carefully removed and replaced with 200 µL of DRAQ7-containing medium (150 nM). DRAQ7 enters cells with compromised plasma membranes and was used for real-time detection of cell death. Cells were imaged at 10× magnification using an IncuCyte S3 Live-Cell Analysis System (Essen Bioscience), with brightfield and red fluorescence channels. Images were acquired every 60 min for 48 h. DRAQ7 signal was quantified as total integrated red fluorescence intensity. At least three independent replicates were performed for each condition, and the standard error was calculated from the respective replicates. In parallel, 0.5 × 106 cells were seeded per well in 6-well plates and transfected with the same master mix to confirm protein expression by Western blot analysis.

### Crystallization, data collection and structure solution

Catalytically inactive TeTIR domain (TEP1*, residues 895–1117) was diluted to 6 mg/mL in 20 mM Tris, pH 7.5, 150 mM NaCl, and 1 mM NAD^+^ and crystallized using sitting-drop vapor diffusion with commercially available sparse-matrix screens. 96-well crystallization plates containing 30 µL of the respective screening conditions were set up with 6 mg/mL protein mixed with the reservoir solution at ratios of 1:2, 1:1, and 2:1 in 300-nL drops. Initial crystals appeared in Crystal HT F8 (Hampton Research; 0.1 M MES monohydrate, pH 6.5, and 1.6 M magnesium sulfate heptahydrate) and MIDAS HT D7 (Molecular Dimensions; 20% (w/v) Jeffamine ED-2003 and 0.1 M HEPES-NaOH, pH 6.5) at 20°C. Crystals harvested from Crystal HT F8 were cryoprotected using 50% saturated sucrose solution in mother liquor, while crystals from the MIDAS condition were frozen directly.

Data from these crystals were collected at beamline P13 operated by EMBL Hamburg at the PETRA III storage ring (DESY, Hamburg, Germany). All datasets were processed using XDS ^51^, and the structures were solved by molecular replacement using PHASER implemented in the Phenix package ^52, 53, 54^ with a model predicted by AlphaFold2 as a search model. The initial solution was automatically rebuilt using Phenix.AutoBuild^55^. As both crystal conditions resulted in nearly identical space groups and virtually identical structures, only the Crystal F8 dataset was further analyzed. The structure was refined using iterative cycles of phenix.refine and REFMAC5 together with assisted model-building using Coot ^56, 57^. For structural analysis and figure generation, UCSF ChimeraX ^58^ and PyMOL Molecular Graphics System, Version 2.1 ^49^ were used.

## Acknowledgment

We thank Claudia Poschner and Shuhua Chen-Winkler for their help. This work was funded by the DFG SFB1403-B06 (Grant 414786233 to KH). Crystals were grown using equipment of the Cologne Crystallization facility (C_2_*f*, http://C2f.uni-koeln.de), which is supported by DFG grant INST 216/949-1 FUGG. Synchrotron data were collected at beamline P13 ^59^ operated by EMBL Hamburg at the PETRA III storage ring (DESY, Hamburg, Germany). We would like to thank David von Stetten for the assistance in using the beamline.

## Author contributions

J.H, N.O. and P.P. performed the biochemical and cell-biological experiments. R.I. performed molecular dynamics and docking experiments, J.J. performed LC-MS experiments, J.G. collected the X-ray data and solved the structure, U.B. supervised crystallography, T.H. and K.H. supervised the biochemical experiments, K.H. conceived the project and contributed bioinformatical analyses. All authors contributed to writing the manuscript.

## Conflict of interest

The authors declare no competing interests.

## Figure Legends

Figure S1: AlphaFold-2 predicted domain organization of human TeTIR proteins.

Predicted aligned error (PAE) plots for full-length human TEP1, NWD1, and NWD2 generated with AlphaFold-2. Amino acid positions are shown on both axes. Low PAE values (green) indicate high confidence in the relative positioning of the corresponding regions. Domain boundaries are indicated by manual annotations based on the predicted domain organization. The labels WD_7_ and WD_5_ denote predicted arrangements of seven or five WD40 repeats, respectively.

**Figure S2: Oligomeric state and NADase activity of TeTIR domains**.

**(A)** Size-exclusion chromatography of human TEP1-TeTIR. The elution profile is shown together with three molecular-weight standards; the apparent molecular mass of TEP1-TeTIR is consistent with the predicted monomeric molecular mass of 25.6 kDa. **(B)** NAM generation in NADase assays with human TEP1-TeTIR and three prokaryotic TeTIR domains from *Arthrobacter sp*. (ArTEP1), *Bacillus cereus* (BcTEP1), and *Methanobacterium subterraneum* (MsTEP1). **(C)** NAM generation by human TEP1-TeTIR, as compared to NWD2 and NPHP3 TeTIR domains, which are hardly active or inactive, respectively.

**Figure S3: Comparison of the TEP1-TeTIR crystal structure and AlphaFold 3 model**. Superposition of the TEP1-TeTIR E1008A crystal structure (green) and the corresponding AlphaFold 3 model (cyan). The structures are highly similar overall, with the main difference occurring in the RWG-containing loop. The region enclosed by the black rectangle is shown enlarged, highlighting the different positions of R942 and W943 in the crystal structure and model. R942 and W943 are shown as sticks and labeled; the rest of the structure is shown in cartoon representation.

**Figure S4: Additional structural analyses of TEP1-TeTIR. (A)** Per-residue root-mean-square fluctuation (RMSF) over the 300 ns MD trajectories shown in Figure 4. **(B)** Relative solvent accessibility of residue 1008 over the same trajectories. For both panels, the original crystal structure (E1008A), the crystal structure with E1008 restored in silico, and the open-pocket structure subsequently re-mutated to E1008A are shown in orange, blue, and green, respectively. **(C)** Comparison of four models of NAD+ binding to TEP1-TeTIR. The upper panels show NAD+-bound models generated by AlphaFold 3 (AF3) and Protenix. The lower panels show two alternative docking models, in which R976 or R942, respectively, is positioned to recognize the NAD^+^ phosphates. All models are shown at the same scale and orientation, with the protein in grey cartoon representation, NAD^+^ in red sticks, and residues contacting NAD^+^ shown as grey sticks.

**Figure S5: Additional analysis of cellular NAD depletion and cell death upon TeTIR expression. (A)** Cellular NAD(H) levels measured 24 h after transfection with human TEP1, TEP1*, SARM1, SARM1*, and the indicated human TEP1 mutants. Mutant names are shown below the corresponding bars. **(B)** Time course of DRAQ7 incorporation over 48 h following transfection with the indicated TeTIR constructs, measured by Incucyte imaging. Murine mTEP1 (dark blue) and hTEP1 (middle blue) show high activity, while the inactive hTEP1* is inactive. hNWD2 (orange) and mNWD2 (light orange) show weak activity, while TTC41 (light pink) and NPHP3 (light green) are not active beyond control level (grey).

## Notes

### Competing Interest Statement

The authors have declared no competing interest.

