## Supplementary material for "TeTIR domains define a new class of animal TIR-like NADase effectors in STAND proteins": Suppl. Fig.

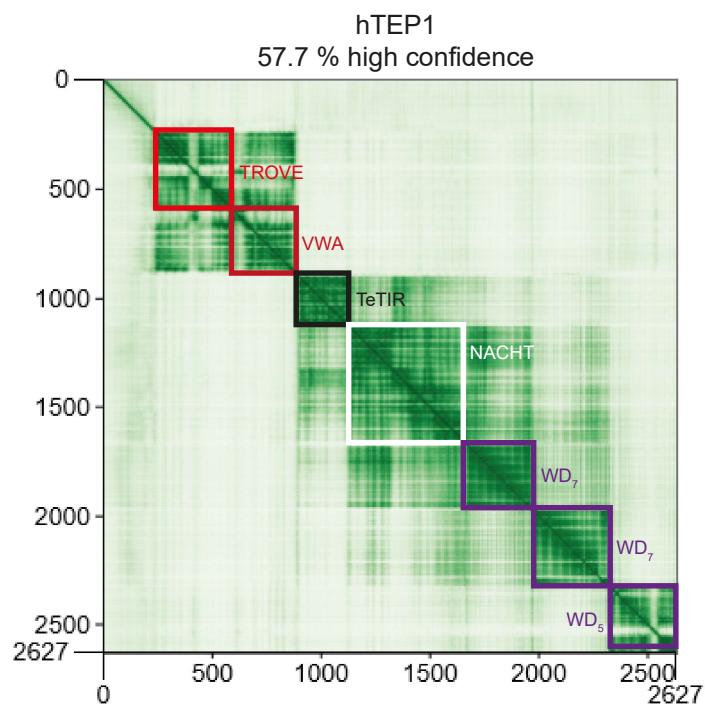

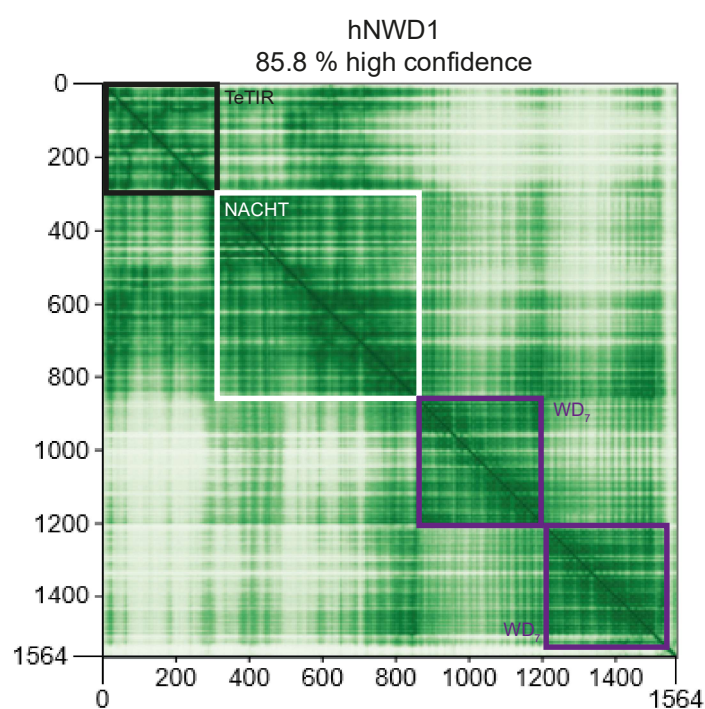

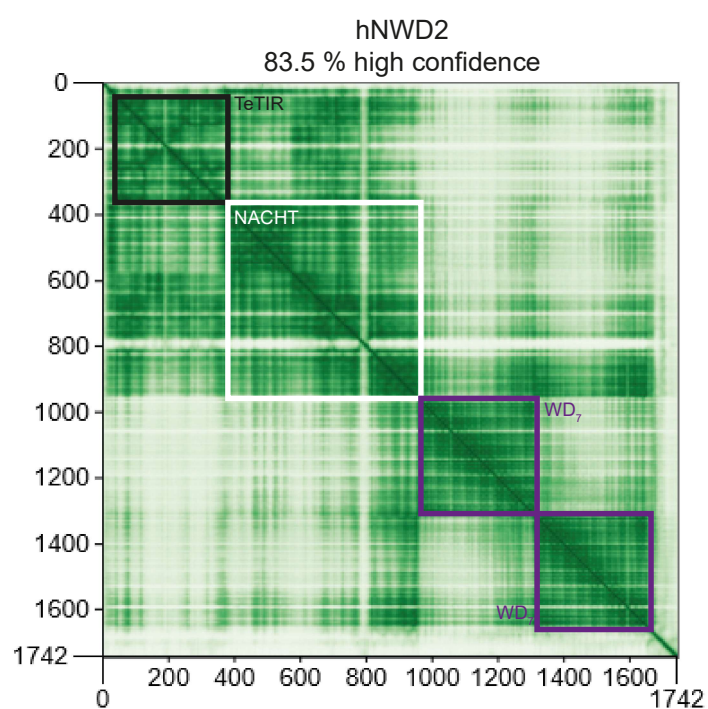

**Figure S1:** AlphaFold-2 predicted domain organization of human TeTIR proteins. Predicted aligned error (PAE) plots for full-length human TEP1, NWD1, and NWD2 generated with AlphaFold-2. Amino acid positions are shown on both axes. Low PAE values (green) indicate high confidence in the relative positioning of the corresponding regions. Domain boundaries are indicated by manual annotations based on the predicted domain organization. The labels WD7 and WD5 denote predicted arrangements of seven or five WD40 repeats, respectively.

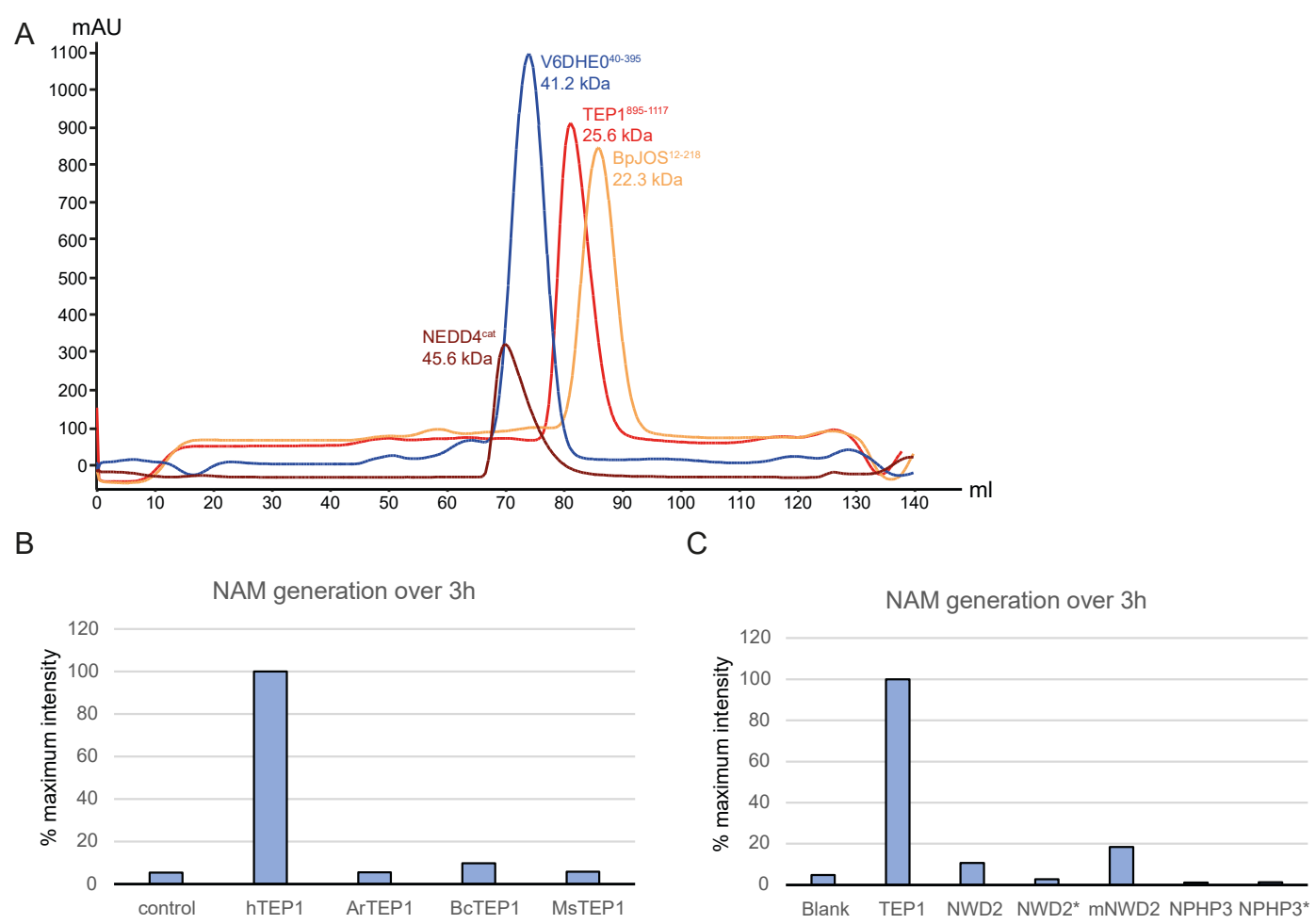

**Figure S2:** Oligomeric state and NADase activity of TeTIR domains. (A) Size-exclusion chromatography of human TEP1-TeTIR. The elution profile is shown together with three molecular-weight standards; the apparent molecular mass of TEP1-TeTIR is consistent with the predicted monomeric molecular mass of 25.6 kDa. (B) NAM generation in NADase assays with human TEP1-TeTIR and three prokaryotic TeTIR domains from *Arthrobacter* sp. (ArTEP1), *Bacillus cereus* (BcTEP1), and *Methanobacterium subterraneum* (MsTEP1). (C) NAM generation by human TEP1-TeTIR, as compared to NWD2 and NPHP3 TeTIR domains, which are hardly active or inactive, respectively.

A

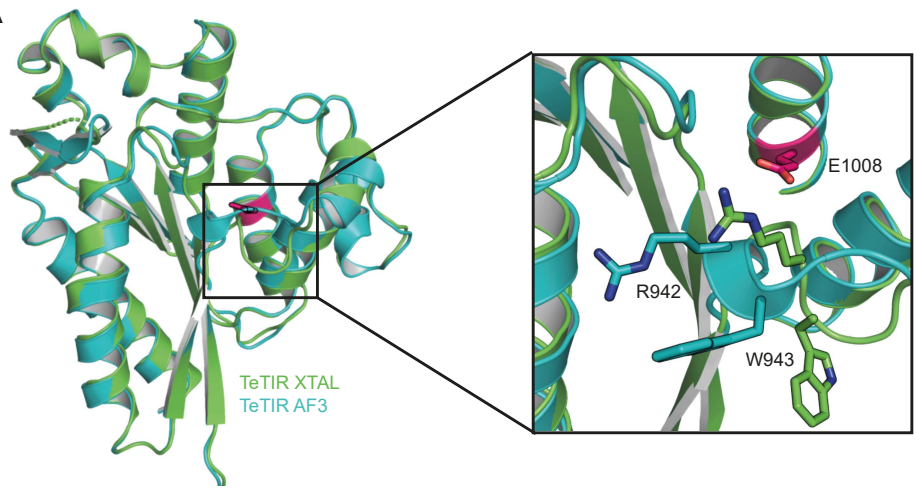

Figure S3: Comparison of the TEP1-TeTIR crystal structure and AlphaFold 3 model. Superposition of the TEP1-TeTIR E1008A crystal structure (green) and the corresponding AlphaFold 3 model (cyan). The structures are highly similar overall, with the main difference occurring in the RWG-containing loop. The region enclosed by the black rectangle is shown enlarged, highlighting the different positions of R942 and W943 in the crystal structure and model. R942 and W943 are shown as sticks and labeled; the rest of the structure is shown in cartoon representation.

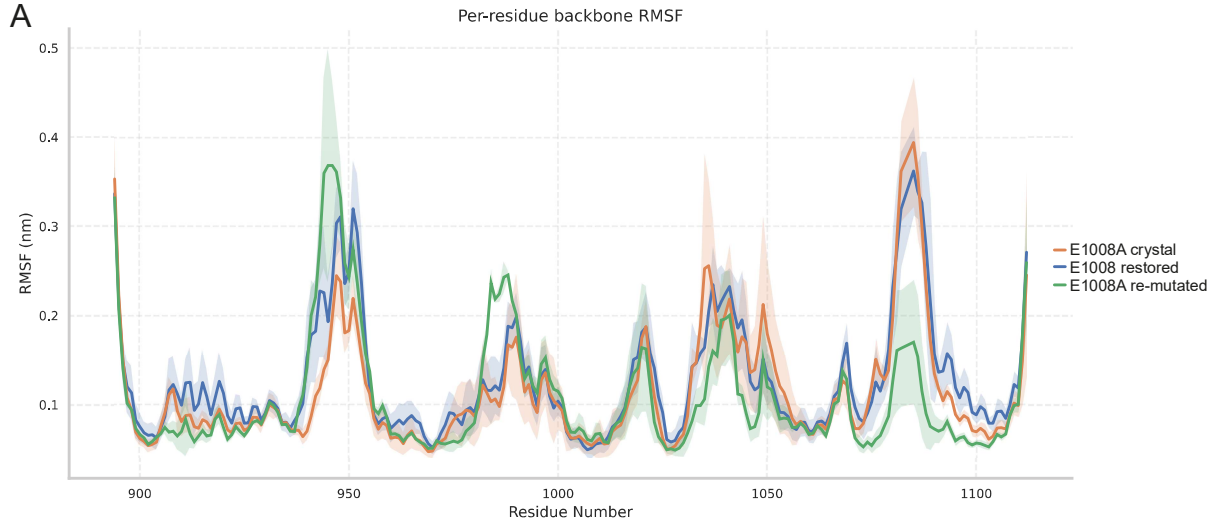

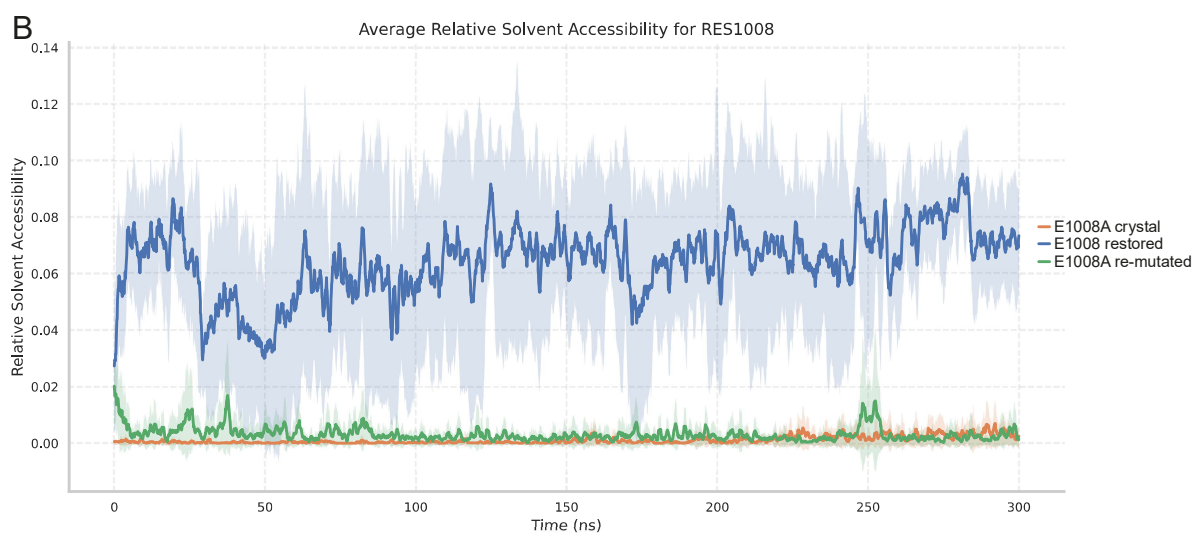

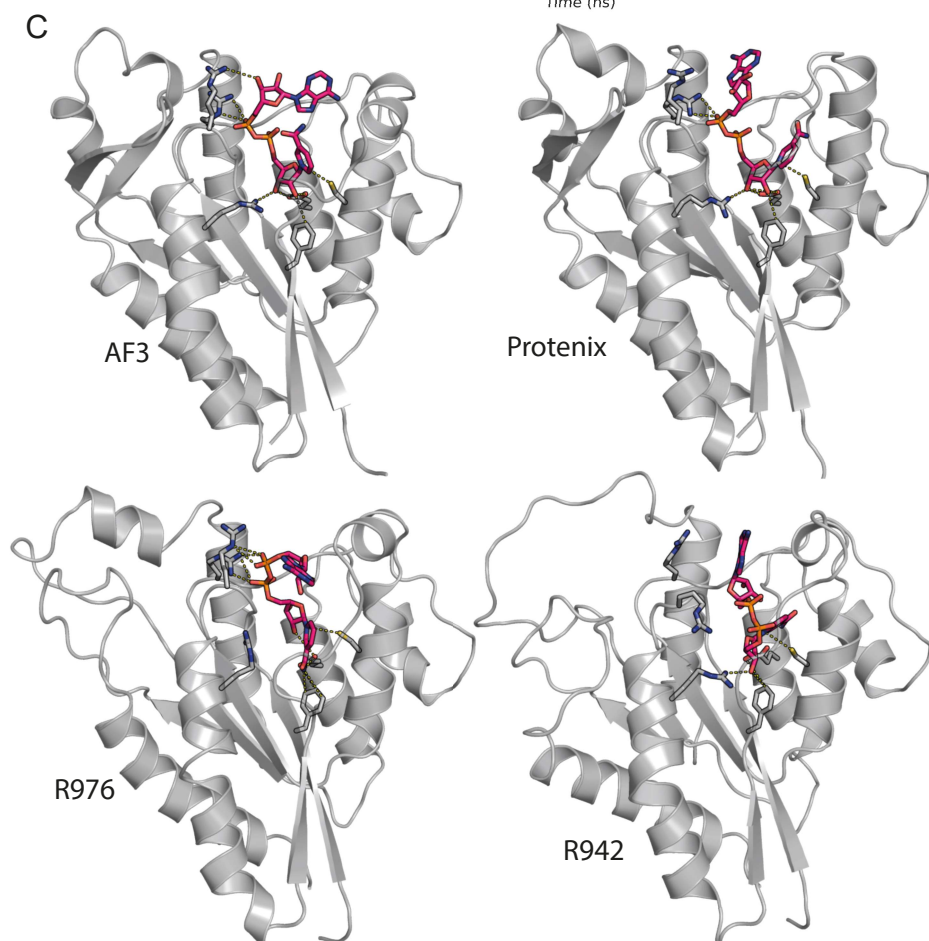

**Figure S4:** Additional structural analyses of TEP1-TeTIR. (A) Per-residue root-mean-square fluctuation (RMSF) over the 300 ns MD trajectories shown in Figure 4. (B) Relative solvent accessibility of residue 1008 over the same trajectories. For both panels, the original crystal structure (E1008A), the crystal structure with E1008 restored in silico, and the open-pocket structure subsequently re-mutated to E1008A are shown in orange, blue, and green, respectively. (C) Comparison of four models of NAD<sup>+</sup> binding to TEP1-TeTIR. The upper panels show NAD<sup>+</sup>-bound models generated by AlphaFold 3 (AF3) and Protenix. The lower panels show two alternative docking models, in which R976 or R942, respectively, is positioned to recognize the NAD<sup>+</sup> phosphates. All models are shown at the same scale and orientation, with the protein in grey cartoon representation, NAD<sup>+</sup> in red sticks, and residues contacting NAD<sup>+</sup> shown as grey sticks.

A

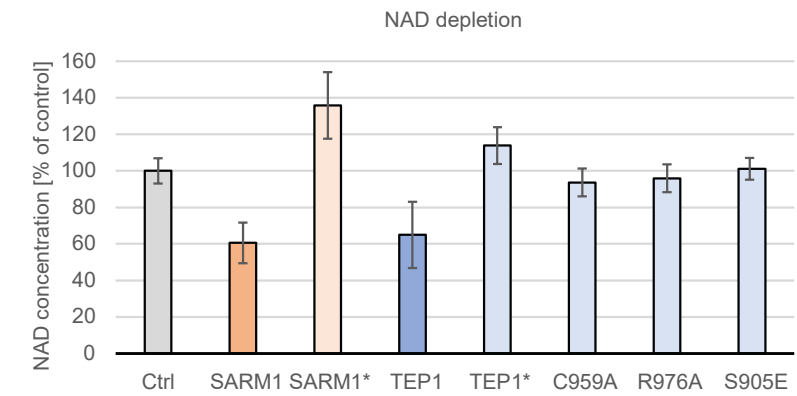

B

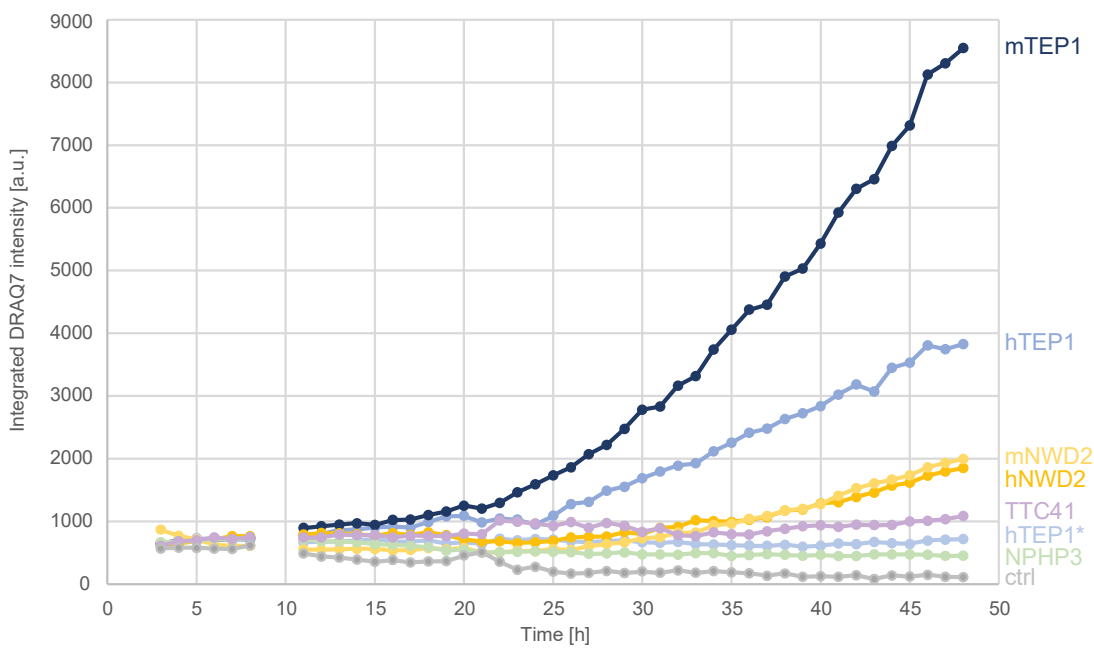

**Figure S5:** Additional analysis of cellular NAD depletion and cell death upon TeTIR expression. (A) Cellular NAD(H) levels measured 24 h after transfection with human TEP1, TEP1\*, SARM1, SARM1\*, and the indicated human TEP1 mutants. Mutant names are shown below the corresponding bars. (B) Time course of DRAQ7 incorporation over 48 h following transfection with the indicated TeTIR constructs, measured by Incucyte imaging. Murine mTEP1 (dark blue) and hTEP1 (middle blue) show high activity, while the inactive hTEP1\* is inactive. hNWD2 (orange) and mNWD2 (light orange) show weak activity, while TTC41 (light pink) and NPHP3 (light green) are not active beyond control level (grey).
