## Supplementary Table 1 for "TeTIR domains define a new class of animal TIR-like NADase effectors in STAND proteins"

| **Name** | **Organism** | **Uniprot** | **Vector** | **Source** | **Protein sequence** |
| --- | --- | --- | --- | --- | --- |
| mTEP1 – TeTIR  902-1126 | Mus musculus | [P97499](https://www.uniprot.org/uniprotkb/P97499/entry) | POPINS | This study | QHGWRNIRLFISSTFRDMHGERDLLMRSVLPALQARVFPHRISLHAIDLRWGITEEETRRNRQLEVCLGEVENSQLFVGILGSRYGYIPPSYDLPDHPHFHWTHEYPSGRSVTEMEVMQFLNRGQRSQPSAQALIYFRDPDFLSSVPDAWKPDFISESEEAAHRVSELKRYLHEQKEVTCRSYSCEWGGVAAGRPYTGGLEEFGQLVLQDVWSMIQKQHLQPGAQ |
| mTEP1 – TeTIR  902-1126 | Mus musculus | [P97499](https://www.uniprot.org/uniprotkb/P97499/entry) | pcDNA5/FRT/TO | This study |  |
| mTEP1 E1017A – TeTIR  902-1126 | Mus musculus |  | POPINS | This study | QHGWRNIRLFISSTFRDMHGERDLLMRSVLPALQARVFPHRISLHAIDLRWGITEEETRRNRQLEVCLGEVENSQLFVGILGSRYGYIPPSYDLPDHPHFHWTHEYPSGRSVTEM**A**VMQFLNRGQRSQPSAQALIYFRDPDFLSSVPDAWKPDFISESEEAAHRVSELKRYLHEQKEVTCRSYSCEWGGVAAGRPYTGGLEEFGQLVLQDVWSMIQKQHLQPGAQ |
| mTEP1 E1017A – TeTIR  902-1126 | Mus musculus |  | pcDNA5/FRT/TO | This study |  |
| TEP1 – TeTIR  895-1117 | Homo sapiens | Q99973 | pcDNA5/FRT/TO | This study | GWRSIRLFISSTFRDMHGERDLLLRSVLPALQARAAPHRISLHGIDLRWGVTEEETRRNRQLEVCLGEVENAQLFVGILGSRYGYIPPSYNLPDHPHFHWAQQYPSGRSVTEMEVMQFLNRNQRLQPSAQALIYFRDSSFLSSVPDAWKSDFVSESEEAARRISELKSYLSRQKGITCRRYPCEWGGVAAGRPYVGGLEEFGQLVLQDVWNMIQKLYLQPGAL |
| TEP1 – TeTIR  895-1117 | Homo sapiens | Q99973 | POPINS | This study |  |
| TEP1 E1008A – TeTIR  895-1117, | Homo sapiens |  | pcDNA5/FRT/TO | This study | GWRSIRLFISSTFRDMHGERDLLLRSVLPALQARAAPHRISLHGIDLRWGVTEEETRRNRQLEVCLGEVENAQLFVGILGSRYGYIPPSYNLPDHPHFHWAQQYPSGRSVTEM**A**VMQFLNRNQRLQPSAQALIYFRDSSFLSSVPDAWKSDFVSESEEAARRISELKSYLSRQKGITCRRYPCEWGGVAAGRPYVGGLEEFGQLVLQDVWNMIQKLYLQPGAL |
| TEP1 E1008A – TeTIR  895-1117, | Homo sapiens |  | POPINS | This study |  |
| TEPE1008Q – TeTIR  895-1117, | Homo sapiens |  | pcDNA5/FRT/TO | This study | GWRSIRLFISSTFRDMHGERDLLLRSVLPALQARAAPHRISLHGIDLRWGVTEEETRRNRQLEVCLGEVENAQLFVGILGSRYGYIPPSYNLPDHPHFHWAQQYPSGRSVTEM**Q**VMQFLNRNQRLQPSAQALIYFRDSSFLSSVPDAWKSDFVSESEEAARRISELKSYLSRQKGITCRRYPCEWGGVAAGRPYVGGLEEFGQLVLQDVWNMIQKLYLQPGAL |
| TEPE1008Q – TeTIR  895-1117, | Homo sapiens |  | POPINS | This study |  |
| TEP1F902E – TeTIR  895-1117, | Homo sapiens |  | pcDNA5/FRT/TO | This study | GWRSIRL**A**ISSTFRDMHGERDLLLRSVLPALQARAAPHRISLHGIDLRWGVTEEETRRNRQLEVCLGEVENAQLFVGILGSRYGYIPPSYNLPDHPHFHWAQQYPSGRSVTEMEVMQFLNRNQRLQPSAQALIYFRDSSFLSSVPDAWKSDFVSESEEAARRISELKSYLSRQKGITCRRYPCEWGGVAAGRPYVGGLEEFGQLVLQDVWNMIQKLYLQPGAL |
| TEP1F902E – TeTIR  895-1117, | Homo sapiens |  | POPINS | This study |  |
| TEP1S905E – TeTIR  895-1117, | Homo sapiens |  | pcDNA5/FRT/TO | This study | GWRSIRLFIS**E**TFRDMHGERDLLLRSVLPALQARAAPHRISLHGIDLRWGVTEEETRRNRQLEVCLGEVENAQLFVGILGSRYGYIPPSYNLPDHPHFHWAQQYPSGRSVTEMEVMQFLNRNQRLQPSAQALIYFRDSSFLSSVPDAWKSDFVSESEEAARRISELKSYLSRQKGITCRRYPCEWGGVAAGRPYVGGLEEFGQLVLQDVWNMIQKLYLQPGAL |
| TEP1S905E – TeTIR  895-1117, | Homo sapiens |  | POPINS | This study |  |
| TEP1R908A – TeTIR  895-1117, | Homo sapiens |  | pcDNA5/FRT/TO | This study | GWRSIRLFISSTF**A**DMHGERDLLLRSVLPALQARAAPHRISLHGIDLRWGVTEEETRRNRQLEVCLGEVENAQLFVGILGSRYGYIPPSYNLPDHPHFHWAQQYPSGRSVTEMEVMQFLNRNQRLQPSAQALIYFRDSSFLSSVPDAWKSDFVSESEEAARRISELKSYLSRQKGITCRRYPCEWGGVAAGRPYVGGLEEFGQLVLQDVWNMIQKLYLQPGAL |
| TEP1R908A – TeTIR  895-1117, | Homo sapiens |  | POPINS | This study |  |
| TEP1R914A – TeTIR  895-1117, | Homo sapiens |  | pcDNA5/FRT/TO | This study | GWRSIRLFISSTFRDMHGE**A**DLLLRSVLPALQARAAPHRISLHGIDLRWGVTEEETRRNRQLEVCLGEVENAQLFVGILGSRYGYIPPSYNLPDHPHFHWAQQYPSGRSVTEMEVMQFLNRNQRLQPSAQALIYFRDSSFLSSVPDAWKSDFVSESEEAARRISELKSYLSRQKGITCRRYPCEWGGVAAGRPYVGGLEEFGQLVLQDVWNMIQKLYLQPGAL |
| TEP1R914A – TeTIR  895-1117, | Homo sapiens |  | POPINS | This study |  |
| TEP1R942A – TeTIR  895-1117, | Homo sapiens |  | pcDNA5/FRT/TO | This study | GWRSIRLFISSTFRDMHGERDLLLRSVLPALQARAAPHRISLHGIDL**A**WGVTEEETRRNRQLEVCLGEVENAQLFVGILGSRYGYIPPSYNLPDHPHFHWAQQYPSGRSVTEMEVMQFLNRNQRLQPSAQALIYFRDSSFLSSVPDAWKSDFVSESEEAARRISELKSYLSRQKGITCRRYPCEWGGVAAGRPYVGGLEEFGQLVLQDVWNMIQKLYLQPGAL |
| TEP1R942A – TeTIR  895-1117, | Homo sapiens |  | POPINS | This study |  |
| TEP1W943A – TeTIR  895-1117, | Homo sapiens |  | pcDNA5/FRT/TO | This study | GWRSIRLFISSTFRDMHGERDLLLRSVLPALQARAAPHRISLHGIDLR**A**GVTEEETRRNRQLEVCLGEVENAQLFVGILGSRYGYIPPSYNLPDHPHFHWAQQYPSGRSVTEMEVMQFLNRNQRLQPSAQALIYFRDSSFLSSVPDAWKSDFVSESEEAARRISELKSYLSRQKGITCRRYPCEWGGVAAGRPYVGGLEEFGQLVLQDVWNMIQKLYLQPGAL |
| TEP1W943A – TeTIR  895-1117, | Homo sapiens |  | POPINS | This study |  |
| TEP1V945A – TeTIR  895-1117, | Homo sapiens |  | pcDNA5/FRT/TO | This study | GWRSIRLFISSTFRDMHGERDLLLRSVLPALQARAAPHRISLHGIDLRWG**A**TEEETRRNRQLEVCLGEVENAQLFVGILGSRYGYIPPSYNLPDHPHFHWAQQYPSGRSVTEMEVMQFLNRNQRLQPSAQALIYFRDSSFLSSVPDAWKSDFVSESEEAARRISELKSYLSRQKGITCRRYPCEWGGVAAGRPYVGGLEEFGQLVLQDVWNMIQKLYLQPGAL |
| TEP1V945A – TeTIR  895-1117, | Homo sapiens |  | POPINS | This study |  |
| TEP1C959A – TeTIR  895-1117, | Homo sapiens |  | pcDNA5/FRT/TO | This study | GWRSIRLFISSTFRDMHGERDLLLRSVLPALQARAAPHRISLHGIDLRWGVTEEETRRNRQLEV**A**LGEVENAQLFVGILGSRYGYIPPSYNLPDHPHFHWAQQYPSGRSVTEMEVMQFLNRNQRLQPSAQALIYFRDSSFLSSVPDAWKSDFVSESEEAARRISELKSYLSRQKGITCRRYPCEWGGVAAGRPYVGGLEEFGQLVLQDVWNMIQKLYLQPGAL |
| TEP1C959A – TeTIR  895-1117, | Homo sapiens |  | POPINS | This study |  |
| TEP1C959A/V945A – TeTIR  895-1117, | Homo sapiens |  | pcDNA5/FRT/TO | This study | GWRSIRLFISSTFRDMHGERDLLLRSVLPALQARAAPHRISLHGIDLRWG**A**TEEETRRNRQLEV**A**LGEVENAQLFVGILGSRYGYIPPSYNLPDHPHFHWAQQYPSGRSVTEMEVMQFLNRNQRLQPSAQALIYFRDSSFLSSVPDAWKSDFVSESEEAARRISELKSYLSRQKGITCRRYPCEWGGVAAGRPYVGGLEEFGQLVLQDVWNMIQKLYLQPGAL |
| TEP1C959A/V945A – TeTIR  895-1117, | Homo sapiens |  | POPINS | This study |  |
| TEP1R976A – TeTIR  895-1117, | Homo sapiens |  | pcDNA5/FRT/TO | This study | GWRSIRLFISSTFRDMHGERDLLLRSVLPALQARAAPHRISLHGIDLRWGVTEEETRRNRQLEVCLGEVENAQLFVGILGS**A**YGYIPPSYNLPDHPHFHWAQQYPSGRSVTEMEVMQFLNRNQRLQPSAQALIYFRDSSFLSSVPDAWKSDFVSESEEAARRISELKSYLSRQKGITCRRYPCEWGGVAAGRPYVGGLEEFGQLVLQDVWNMIQKLYLQPGAL |
| TEP1R976A – TeTIR  895-1117 | Homo sapiens |  | POPINS | This study |  |
| TEP1Y979A – TeTIR  895-1117 | Homo sapiens |  | pcDNA5/FRT/TO | This study | GWRSIRLFISSTFRDMHGERDLLLRSVLPALQARAAPHRISLHGIDLRWGVTEEETRRNRQLEVCLGEVENAQLFVGILGSRYG**A**IPPSYNLPDHPHFHWAQQYPSGRSVTEMEVMQFLNRNQRLQPSAQALIYFRDSSFLSSVPDAWKSDFVSESEEAARRISELKSYLSRQKGITCRRYPCEWGGVAAGRPYVGGLEEFGQLVLQDVWNMIQKLYLQPGAL |
| TEP1Y979A – TeTIR  895-1117 | Homo sapiens |  | POPINS | This study |  |
| drTEP1 – TeTIR  896-1113 | Danio rerio | A0A8M3AXG0 | pcDNA5/FRT/TO | This study | KFRWKGVRVFISSTFRDMHAERDVLVRSVFPELRRRAAPHYLYLQELELRWGVTEEESNRAAELCLSEVCRSQLLLGILGERYGLVPPRPILPDLPQYNWLKSASDGLSITEMEIRQFQALHPDSAQSRMFFYFRSPHLARSVPAAWRTDFVAESKEAEAKMNSLKTWIQSSEFKVTENYPCEWGGVMDGKPYVKGLEEFGRAALEDIWEAVQQLFVE |
| drTEP1 – TeTIR  896-1113 | Danio rerio | A0A8M3AXG0 | POPINS | This study |  |
| drTEP1 E1009A – TeTIR  896-1113 | Danio rerio |  | pcDNA5/FRT/TO | This study | KFRWKGVRVFISSTFRDMHAERDVLVRSVFPELRRRAAPHYLYLQELELRWGVTEEESNRAAELCLSEVCRSQLLLGILGERYGLVPPRPILPDLPQYNWLKSASDGLSITEM**A**IRQFQALHPDSAQSRMFFYFRSPHLARSVPAAWRTDFVAESKEAEAKMNSLKTWIQSSEFKVTENYPCEWGGVMDGKPYVKGLEEFGRAALEDIWEAVQQLFVE |
| drTEP1 E1009A – TeTIR  896-1113 | Danio rerio |  | POPINS | This study |  |
| aqTEP1 – TeTIR  1012-1230 | Amphimedon queenslandica | A0AANOIPT6 | pcDNA5/FRT/TO | This study | QTKWRTARIFISSTFRDMHGERDLLTRYVFPELRRRAHDLRVYLYEVDLRWGVTEEDTKHHKAIEICMTEISKSNYFIGILGQRYGWCPGEYSVPDSEEYDWIRDYPKGRSITELEMYHGALADPDKVAGKAFFYFRDKGVMDEIPTKYIGHFKDESFENEEKIESLKSKIRTSGIEVFDNYPSHWMGVVQNKPMLSSLEDFGKRVINNLWNTIQRDYP |
| aqTEP1 – TeTIR  1012-1230 | Amphimedon queenslandica | A0AANOIPT6 | POPINS | This study |  |
| aqTEP1 E1127A – TeTIR  1012-1230 | Amphimedon queenslandica |  | pcDNA5/FRT/TO | This study | QTKWRTARIFISSTFRDMHGERDLLTRYVFPELRRRAHDLRVYLYEVDLRWGVTEEDTKHHKAIEICMTEISKSNYFIGILGQRYGWCPGEYSVPDSEEYDWIRDYPKGRSITEL**A**MYHGALADPDKVAGKAFFYFRDKGVMDEIPTKYIGHFKDESFENEEKIESLKSKIRTSGIEVFDNYPSHWMGVVQNKPMLSSLEDFGKRVINNLWNTIQRDYP |
| aqTEP1 E1127A – TeTIR  1012-1230 | Amphimedon queenslandica |  | POPINS | This study |  |
| ddTEP1 – TeTIR  696-919 | *Dictyostelium discoideum* | Q54CB5 | pcDNA5/FRT/TO | This study | KPWKNAKVFISSTFLDMQGERDLLVKTIFPELRARCLKNRIHLTEIDLRWGITEEDALKNRSVDLCLEEVDRCRPFFISLLGQRYGWVPKKEQIPQDSKYDWIRELPTQRSITELEVLYASFQGKKKPTTNSLVYFRDPSFSQEVPKQFRDQFECEDRTSQSKLEQLKHKITQSKSQHLSHYTYSAKWGGVNSEGIPVSTGLDELCERIQTDLWDNICKVFNLN |
| ddTEP1 – TeTIR  696-919 | *Dictyostelium discoideum* | Q54CB5 | POPINS | This study |  |
| ddTEP1 E811A – TeTIR  696-919 | *Dictyostelium discoideum* |  | pcDNA5/FRT/TO | This study | KPWKNAKVFISSTFLDMQGERDLLVKTIFPELRARCLKNRIHLTEIDLRWGITEEDALKNRSVDLCLEEVDRCRPFFISLLGQRYGWVPKKEQIPQDSKYDWIRELPTQRSITEL**A**VLYASFQGKKKPTTNSLVYFRDPSFSQEVPKQFRDQFECEDRTSQSKLEQLKHKITQSKSQHLSHYTYSAKWGGVNSEGIPVSTGLDELCERIQTDLWDNICKVFNLN |
| ddTEP1 E811A – TeTIR  696-919 | *Dictyostelium discoideum* |  | POPINS | This study |  |
| ArTEP1– TeTIR  1-180 | [*Arthrobacter sp.*](https://www.uniprot.org/taxonomy/2292263) | A0A371BSX1 | pcDNA5/FRT/TO | This study | MIDSTAWPRVGLFISSTFDDMHGERDYLVKRVLPKLYQWCEERRLHLVDVDLRWGINREVEVVGSCLQAIDACQPLFVCFLGQRFGRVTAIEVEHALQQGSRVIFYLRDPDYLADLPAELRFVYANGHENEMQHLRDLAREEARTYSAHWNPDTGRLTEFDFGDRLLADLQSAIASAYPD |
| ArTEP1– TeTIR  1-180 | [*Arthrobacter sp.*](https://www.uniprot.org/taxonomy/2292263) | A0A371BSX1 | POPINS | This study |  |
| ArTEP1 E92A–  TeTIR 1-180 | [*Arthrobacter sp.*](https://www.uniprot.org/taxonomy/2292263) |  | pcDNA5/FRT/TO | This study | MIDSTAWPRVGLFISSTFDDMHGERDYLVKRVLPKLYQWCEERRLHLVDVDLRWGINREVEVVGSCLQAIDACQPLFVCFLGQRFGRVTAI**A**VEHALQQGSRVIFYLRDPDYLADLPAELRFVYANGHENEMQHLRDLAREEARTYSAHWNPDTGRLTEFDFGDRLLADLQSAIASAYPD |
| ArTEP1 E92A–  TeTIR 1-180 | [*Arthrobacter sp.*](https://www.uniprot.org/taxonomy/2292263) |  | POPINS | This study |  |
| SsTEP1 – TeTIR  14-307 | *Streptomyces* | A0A0MWUZ4 | pcDNA5/FRT/TO | This study | VAVNTFRVFVSSTFGDLAEERAALHERVFPRLRALCSEADAEFQAIDLRWGVSEEAAIEQRVAEICLGEIDRCRQITRRPNFLVLLGQRYGARPAPPRIDAAMFRRLCARLTAEQGRLVNRWYVEDRNHVPAVFVLQPRRGGYRDDALWRPTERALSEALTAAALQEPAEREAVALLRTSITEQEILRGIPENGPEHGAAFAFQRTILGLPEDASAAPYRDFLDERPDTEAAAAQQLLRDRLSAMGAARLRLTDYRAAWDYELGRPTTGHIDRLCEDVYASLAEVIRDELQAAE |
| SsTEP1 – TeTIR  14-307 | *Streptomyces* | A0A0MWUZ4 | POPINS | This study |  |
| SsTEP1 E198A - TeTIR  14-307 | *Streptomyces* |  | pcDNA5/FRT/TO | This study | VAVNTFRVFVSSTFGDLAEERAALHERVFPRLRALCSEADAEFQAIDLRWGVSEEAAIEQRVAEICLGEIDRCRQITRRPNFLVLLGQRYGARPAPPRIDAAMFRRLCARLTAEQGRLVNRWYVEDRNHVPAVFVLQPRRGGYRDDALWRPTERALSEALTAAALQEPAEREAVALLRTSITEQ**A**ILRGIPENGPEHGAAFAFQRTILGLPEDASAAPYRDFLDERPDTEAAAAQQLLRDRLSAMGAARLRLTDYRAAWDYELGRPTTGHIDRLCEDVYASLAEVIRDELQAAE |
| SsTEP1 E198A – TeTIR  14-307 | *Streptomyces* |  | POPINS | This study |  |
| MsTEP1– TeTIR  1-214 | *Methanobacterium subterraneum* | A0A2H4VP27 | pcDNA5/FRT/TO | This study | KNSWKTVKIFISSTFKDMQAERDHLVRFVFPRLREDLIKYKIYLIDVDLRWGITSDQDAYDVCMDEIDNCHPYFMCMLGGRYGWIPPEKKISITESEIYYGALDKLDIPTFRFFYLRDAKITNSIPAEFNDDYKELKDSESEKQLKYLKEKIMNPEIKGKVLIKPDEVECKSLSYFIYPCKWDNNLKRIVDLESFGNQVYNDIMSSINVELG |
| MsTEP1– TeTIR  1-214 | *Methanobacterium subterraneum* | A0A2H4VP27 | POPINS | This study |  |
| MsTEP1 E98A – TeTIR  1-214 | *Methanobacterium subterraneum* |  | pcDNA5/FRT/TO | This study | KNSWKTVKIFISSTFKDMQAERDHLVRFVFPRLREDLIKYKIYLIDVDLRWGITSDQDAYDVCMDEIDNCHPYFMCMLGGRYGWIPPEKKISITES**A**IYYGALDKLDIPTFRFFYLRDAKITNSIPAEFNDDYKELKDSESEKQLKYLKEKIMNPEIKGKVLIKPDEVECKSLSYFIYPCKWDNNLKRIVDLESFGNQVYNDIMSSINVELG |
| MsTEP1 E98A – TeTIR  1-214 | *Methanobacterium subterraneum* |  | POPINS | This study |  |
| BcTEP1 – TeTIR  1-210 | [*Bacillus cereus*](https://www.uniprot.org/taxonomy/1396) | A0A2B0L8K0 | pcDNA5/FRT/TO | This study | MWETIRIFVSSTFIDMSEERDWLVQHVFPKLRKKCEERKLRLVDVDLRWGIPEGKITDENLADICLEEIENCNPFFVCLLGSRYGTLAKVNEKRKAELMLQDEQHSITALEIYHRLRKMQEKKGGVLFYFRDVSPIKGVPKDVNSEGDFKKKLAELKDDIYKEVDHTLIKDYSHLYSKDEQSEFMKLNKVAFGNLVFEDLWKQIDEMYPV |
| BcTEP1 – TeTIR  1-210 | [*Bacillus cereus*](https://www.uniprot.org/taxonomy/1396) | A0A2B0L8K0 | POPINS | This study |  |
| BcTEP1 E111A – TeTIR  1-210 | [*Bacillus cereus*](https://www.uniprot.org/taxonomy/1396) |  | pcDNA5/FRT/TO | This study | MWETIRIFVSSTFIDMSEERDWLVQHVFPKLRKKCEERKLRLVDVDLRWGIPEGKITDENLADICLEEIENCNPFFVCLLGSRYGTLAKVNEKRKAELMLQDEQHSITAL**A**IYHRLRKMQEKKGGVLFYFRDVSPIKGVPKDVNSEGDFKKKLAELKDDIYKEVDHTLIKDYSHLYSKDEQSEFMKLNKVAFGNLVFEDLWKQIDEMYPV |
| BcTEP1 E111A – TeTIR  1-210 | [*Bacillus cereus*](https://www.uniprot.org/taxonomy/1396) |  | POPINS | This study |  |
| mNWD2 – TeTIR  1-364 | Homo sapiens | [Q6P5U7](https://www.uniprot.org/uniprotkb/Q6P5U7/entry) | pcDNA5/FRT/TO | This study | WPAGAGTKLPCPRDSALRRAAFSGNLTALPSHLVPAGRSVRVFISANPEDTGAERQALRETVYPKLREFCRENYGLEFQVIDLYWGIEEDEWDSPELQKMRMKLLEECLKTSAGPCFVGLLGEKYGNIRIPGEVEASEFEMILDAAVEAKLETKLLEDWYCRDENSVPAAYYLRPRLEVPRSNKNSTQPSASSEQERPWQEISDEIKTIFKAAVKLLHEQGKMKQSQAKRYLFSAIEDEFDFALGKQTPAFLKKCVCYIRKIANIERFVKIPEMGKYMDITGTDPRIVRDPEAQEKLIKLRDEFIPTIVASSNLRVYTSVTHCDMKLGYSQEIENHYIEGLGKQFYEDMIDIIQATVQQNFDT |
| mNWD2 – TeTIR  1-364 | Homo sapiens | [Q6P5U7](https://www.uniprot.org/uniprotkb/Q6P5U7/entry) | POPINS | This study |  |
| NWD2 – TeTIR  35-364 | Homo sapiens | [Q9ULI1](https://www.uniprot.org/uniprotkb/Q9ULI1/entry) | pcDNA5/FRT/TO | This study | VPAGRSVRVFISANPEDTGAERQALRENVYPKLREFCRENYGLEFQVIDLYWGVEEDEWDSPELQKTRMKLLENCLKTSAGPCFVGLLGEKYGNIRIPGEVEASEFEMILDAAIEAKLETKLLEEWYCRDENSVPAAYYLRPKSEMLRSNRNAMQPSTNAENEKTWQEISDEIKKIFKAAVKLLHEKGKMKHSQAKRYLFSAIEDEFDFALGKQTPAFLKKCVCYIRKIANIERFVKIPEMGKYMDITGTEPRIIRDPEAQEKLIKLRDEFIPTIVASSNLRVYTSVTHCDMKLGYSQEIENHYIEGLGKQFYEDMIDIIQATIQQNFDT |
| NWD2 – TeTIR  35-364 | Homo sapiens | [Q9ULI1](https://www.uniprot.org/uniprotkb/Q9ULI1/entry) | POPINS | This study |  |
| NWD2 E240A – TeTIR  35-364 | Homo sapiens |  | pcDNA5/FRT/TO | This study | VPAGRSVRVFISANPEDTGAERQALRENVYPKLREFCRENYGLEFQVIDLYWGVEEDEWDSPELQKTRMKLLENCLKTSAGPCFVGLLGEKYGNIRIPGEVEASEFEMILDAAIEAKLETKLLEEWYCRDENSVPAAYYLRPKSEMLRSNRNAMQPSTNAENEKTWQEISDEIKKIFKAAVKLLHEKGKMKHSQAKRYLFSAIED**A**FDFALGKQTPAFLKKCVCYIRKIANIERFVKIPEMGKYMDITGTEPRIIRDPEAQEKLIKLRDEFIPTIVASSNLRVYTSVTHCDMKLGYSQEIENHYIEGLGKQFYEDMIDIIQATIQQNFDT |
| NWD2 E240A – TeTIR  35-364 | Homo sapiens |  | POPINS | This study |  |
| SARM1 – TIR  559-700 | Homo sapiens |  | pcDNA5/FRT/TO | This study | GDTPDVFISYRRNSGSQLASLLKVHLQLHGFSVFIDVEKLEAGKFEDKLIQSVMGARNFVLVLSPGALDKCMQDHDCKDWVHKEIVTALSCGKNIVPIIDGFEWPEPQVLPEDMQAVLTFNGIKWSHEYQEATIEKIIRFLQ |
| SARM1 – TIR  559-700 | Homo sapiens |  | POPINS | This study |  |
| SARM1 E642A – TIR  559-700 | Homo sapiens |  | pcDNA5/FRT/TO | This study | GDTPDVFISYRRNSGSQLASLLKVHLQLHGFSVFIDVEKLEAGKFEDKLIQSVMGARNFVLVLSPGALDKCMQDHDCKDWVHK**A**IVTALSCGKNIVPIIDGFEWPEPQVLPEDMQAVLTFNGIKWSHEYQEATIEKIIRFLQ |
| SARM1 E642A – TIR  559-700 | Homo sapiens |  | POPINS | This study |  |
| NPHP3 TeTIR  234-470 | Homo sapiens | Q7Z494 | pcDNA5/FRT/TO | This study | GGALGSEPSIGSMIQLQQSFRGPEFAHSSIDVEGPFANVNRDDWDIAVASLLQVTPLFSHSLWSNTVRCYLIYTDETQPEMDLFLKDYSPKLKRMCETMGYFFHAVYFPIDVENQYLTVRKWEIEKSSLVILFIHLTLPSLLLEDCEEAFLKNPEGKPRLIFHRLEDGKVSSDSVQQLIDQVSNLNKTSKAKIIDHSGDPAEGVYKTYICVEKIIKQDILGFENTDLETKDLGSEDS |
| NPHP3 TeTIR  234-470 | Homo sapiens | Q7Z494 | POPINS | This study |  |
| RPP1 – TIR  1-254 | *Arabidopsis thaliana* | F4J339 | pcDNA5/FRT/TO | This study | MGSVMSLGCSKRKATNQDVDSESRKRRKICSTNDAENCRFIQDESSWKHPWSLCANRVISVAAVALTNFRFQQDNQESNSSSLSLPSPATSVSRNWKHDVFPSFHGADVRRTFLSHIMESFRRKGIDTFIDNNIERSKSIGPELKEAIKGSKIAIVLLSRKYASSSWCLDELAEIMKCRQMVGQIVMTIFYEVDPTDIKKQTGEFGKAFTKTCRGKPKEQVERWRKALEDVATIAGYHSHSWRNEADMIEKIST |
| RPP1 – TIR 1-254 | *Arabidopsis thaliana* | F4J339 | POPINS | This study |  |
| TcpC full length | [Escherichia coli O6:H1](https://www.uniprot.org/taxonomy/199310) | A0A0H2V8B5 | pcDNA5/FRT/TO | This study | MIAYENIEFFICLVNVLGNNMYNILFFIFLSIAIPFLLFLAWKQHLKTKEIRSYLLKEGYNIIFNGEGNSYLAFNISNATFRAGNLTSNDYFQASISYIHDYRWEWKEVEAKKINNIFIIYISNIDFPSQKLFYRNNKSLAEIDWAKLQAIFHQPYEIQNDVMQDNNNTHYDFFISHAKEDKDTFVRPLVDELNRLGVIIWYDEQTLEVGDSLRRNIDLGLRKANYGIVILSHNFLNKKWTQYELDSLINRAVYDDNKIILPIWHNINAQEVSKYSHYLADKMALQTSLYSVKEIARELAEIAYRRR |
| TcpC full length | [Escherichia coli O6:H1](https://www.uniprot.org/taxonomy/199310) | A0A0H2V8B5 | POPINS | This study |  |
| TTC41 TeTIR  20-259 | Mus musculus | Q692V3 | pcDNA5/FRT/TO | This study | RQKPILPYICSTLDFQEERDFLAKSIFPRLNDICSSRGTYFKAVDLRWSAVKAHKSFTSNQFRQYSCLQSQHLKLSLDYVNRCFPFFIGLLGQTYGDFLPDYTPFLLSQVKDFESLSKGKKNLYIAAKNGYPWVLKTPNCSLTEFEIIQAVFRKKSQFQFFYFRTSNSLLRTFNEEEEEEEEKLSSAYLLNEQGKMKVGKLKAKIIGKGLPVRFYRDLEELGDMVWKDWSAVVEKLYPFTT |
| TTC41 TeTIR  20-259 | Mus musculus | Q692V3 | POPINS | This study |  |
